# A systematic cross-modal approach identifies astrocytic VCAM1 as a regulator of hippocampal synapse development

**DOI:** 10.64898/2026.03.17.712320

**Authors:** Cydric Geyskens, Efstathia Kotoula, Kjen Bogaert, Elke Leysen, Joris Vandenbempt, Francisco Pestana, Lars Lefever, Pedro Magalhães, Dan Dascenco, Joris de Wit

## Abstract

Astrocyte-mediated cell-cell interactions at synapses are essential for circuit formation, yet the molecular mechanisms underlying astrocyte-mediated regulation of hippocampal synapse development remain incompletely characterized. Here, we take a systematic cross-modal approach to identify novel astrocytic cell surface proteins (CSPs) at the tripartite synapse. By using single-cell spatial transcriptomics (scST) targeting CSPs identified in a previous hippocampal synaptic dataset, we find 10 potential candidate astrocytic CSPs. Subsequent systematic protein-level profiling using multiple antibody-based assays establishes GPR37L1, HepaCAM, and VCAM1 as astrocytic peri-synaptic CSPs. To gain insight into the molecular context in which these proteins operate at synapses, we map their synaptic interaction partners using affinity purification-mass spectrometry (AP-MS), revealing distinct hippocampal synaptic interactomes for GPR37L1 and VCAM1. We then develop a custom multi-metric image-based synapse analysis pipeline to assess the roles of GPR37L1 and VCAM1 in synaptic development, using CRISPR/Cas9-mediated gene knockout (KO) in the mouse hippocampus. While loss of GPR37L1 does not substantially affect excitatory and inhibitory synapses, VCAM1 loss impairs excitatory hippocampal synapse development. Conversely, addition of recombinant VCAM1 to cultured hippocampal neurons increases the density of excitatory synapses. Together, these results identify astrocytic VCAM1 as a regulator of hippocampal synapse development.

## INTRODUCTION

The brain harbors billions of neurons that are intricately connected through highly specialized intercellular junctions known as synapses. These structures enable information processing within neural circuits that underlie cognition and behavior. The development of synapses is essential for the establishment of functional neuronal circuits^1^. Over the past few decades, foundational studies have shown that neuronal cell surface proteins (CSPs) are key regulators of synaptic development by mediating trans-synaptic cell-cell interactions between pre- and postsynaptic compartments^2,3^.

Astrocytes, the most abundant glial cells in the central nervous system, extend highly elaborate, fine processes that sense, process and modulate neuronal activity and synaptic transmission^4^. Electron microscopy and super-resolution imaging have revealed that peri-synaptic astrocytic processes (PAPs) ensheathe the majority of CNS synapses, positioning astrocytes to detect neurotransmitter release and dynamically regulate synaptic function^5,6^. Once considered primarily homeostatic support cells, astrocytes are now recognized as active and essential participants in synaptic function, giving rise to the concept of the “tripartite synapse”^7^.

Seminal studies demonstrated that astrocytic secreted proteins promote synapse formation and function^8–10^. More recently, contact-dependent mechanisms mediated by astrocytic cell surface proteins have emerged as an additional mode of synaptic development regulation^11,12^. These mechanisms have been characterized predominantly in cortical circuits. In contrast, less is known about astrocyte-mediated signaling in the hippocampus, a structure essential for learning and memory that exhibits highly precise laminar-specific synaptic organization. Notably, evidence suggests that astrocytes are transcriptomically diverse between the cortex and hippocampus^13,14^. Consistent with this idea, only two astrocytic CSPs, Ephrin-B1^15,16^ and CD38^17^, have been implicated in hippocampal synaptic development to date. Thus, the molecular repertoire of astrocytic surface proteins shaping hippocampal circuits remains incompletely defined.

Previously, we characterized the hippocampal CA3 synaptic proteome through fluorescent-activated synaptosome sorting (FASS)^18^. Here, we systematically interrogate this proteomic resource to identify astrocytic CSPs positioned to regulate synapse development. Targeted scST revealed that 10 CSPs from this dataset are potentially expressed by astrocytes. Through rigorous characterization using immunohistochemistry (IHC), single-molecule fluorescent in situ hybridization (smFISH), and synaptogliosome preparations, we established GPR37L1, HepaCAM and VCAM1 as a high-confidence set of astrocytic hippocampal peri-synaptic CSPs. To better understand the molecular environment in which these proteins function at synapses, we identified their interaction partners using affinity purification–mass spectrometry (AP–MS), revealing distinct hippocampal synaptic interactomes for GPR37L1 and VCAM1. Subsequently, we combined a CRISPR/Cas9-mediated gene KO approach with a custom computational image analysis pipeline to assess the impact of GPR37L1 and VCAM1 on hippocampal excitatory and inhibitory synapse development in a lamina-resolved manner, revealing evidence for VCAM1’s role in hippocampal excitatory synapse development. Further experiments in hippocampal neuron cultures suggest a role for VCAM1 in selectively promoting excitatory synapse formation. In contrast, GPR37L1 KO did not robustly affect hippocampal synaptic development *in vivo*. Together, our findings uncover VCAM1 as an astrocytic CSP modulating excitatory synapse development in the hippocampus.

## RESULTS

### Proteome-informed targeted single-cell spatial transcriptomics identifies hippocampal astrocytic CSPs

We previously characterized the cell-surface proteome of isolated hippocampal mossy fiber synapses, which reside in the hippocampal CA3 region, identifying approximately 70 enriched CSPs, several of which have undefined synaptic functions^18^. To explore the cell-type-specific expression of these CSPs in a high-throughput manner, we leveraged a targeted image-based single-cell spatial transcriptomics platform (scST, Molecular Cartography, Resolve Biosciences, Germany). To this end, cryosections of P28 wild-type mice containing the hippocampus, matched in age to those used for the CA3 FASS proteome dataset, were subjected to the Molecular Cartography protocol (**Fig. 1A**). This protocol involves hybridizing transcript-specific probes to target mRNAs, followed by several rounds of colorizing and de-colorizing chemistry to accurately identify individual transcripts through a specific barcode, enabling visualization of millions of individual transcripts with subcellular resolution. We selected the full panel of 75 CSPs identified at mossy fiber synapses together with 20 cell-type-specific markers for neurons (*Rbfox3*, *Syt1*, *Snap25*, *Tubb3*), excitatory neurons (*Slc17a6*, *Slc17a7*), inhibitory neurons (*Gad1*, *Gad2*), astrocytes (*Aldh1l1*, *Aldoc*, *Aqp4*, *Slc1a3*, *Slc1a2*), microglia (*Csfr1*, *C1qa*), oligodendrocytes (*Mog*, *Plp1*) and endothelial cells (*Flt1*, *Pecam1*).

**Fig. 1.**
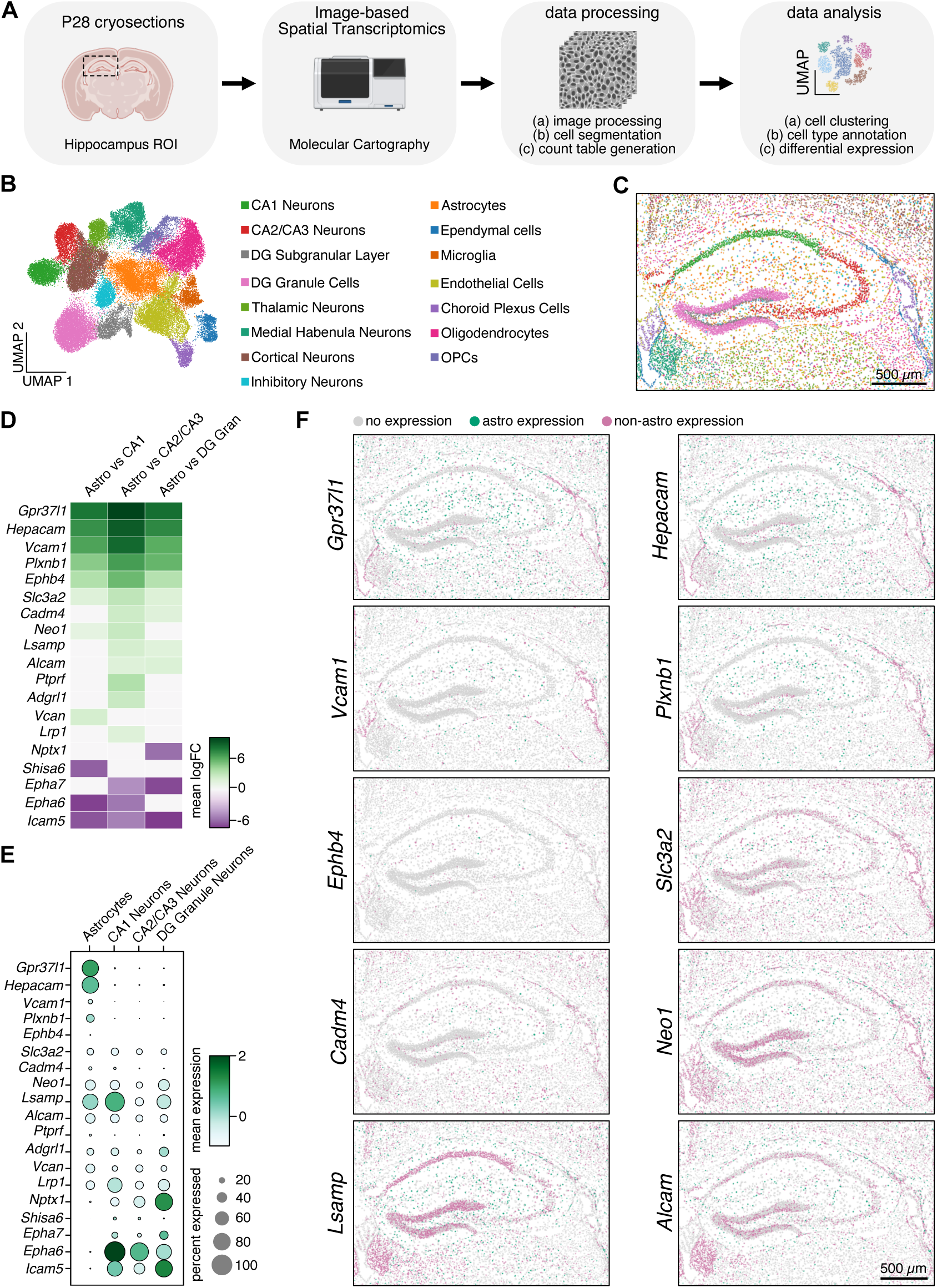
Proteome-informed targeted single-cell spatial transcriptomics identifies hippocampal astrocytic CSPs. (A) Schematic of the workflow used to investigate the hippocampal cell type specificity of the mossy fiber cell-surface synapse proteome. Created with BioRender.com (B) UMAP plot of 15 different clusters annotated with hippocampal cell types (see also Fig S1 for QC). (C) Spatial distribution of the 15 identified cell types in a representative sample (sample 1). (D) Heatmap showing ranked log fold change of three differential expression analyses; astrocytes vs CA1 neurons, astrocytes vs CA2/CA3 neurons, and astrocytes vs DG granule cells. All CSPs enriched in astrocytes are shown, together with top three neuron-enriched CSPs. FDR < 0.05 and mean logFC > 1. (E) Dot plot showing expression of the CSPs shown in (D) across astrocytes, CA1 neurons, CA2/CA3 neurons and DG granule cells. (F) Spatial map of cells indicating CSP expression in astrocytes (green) and non-astrocytes (pink), non-expressing cells are shown in light grey.

We used nuclear DAPI staining to segment the hippocampal data into putative single-cells, followed by batch-corrected scST analysis with extensive quality control (**Fig. 1A** and **S1A-E**). This analysis yielded 15 clusters (**Fig. 1B-C**), with cluster identities confirmed through cell-type-specific markers, differentially expressed genes (DEGs) and their spatial distribution patterns (**Fig. S1F-G**). We could identify the three major hippocampal excitatory neuronal populations: CA1 pyramidal neurons, CA2/CA3 pyramidal neurons and dentate gyrus (DG) granule cells. Despite no inclusion of markers specific to the different cellular layers of the DG, our data resolves distinct subgranular and granular clusters, indicating that they likely express different combinations of synaptic CSPs. Other hippocampus-adjacent neuronal populations included thalamic neurons, cortical neurons and medial habenula neurons as well as inhibitory neurons across the hippocampus and adjacent regions. We could also distinguish endothelial cells, choroid plexus cells and ependymal cells. Glial cell types included astrocytes, microglia, oligodendrocytes and oligodendrocyte precursor cells (OPCs, based on their CSP DEGs and spatial distribution).

As astrocytes display regional heterogeneity within the hippocampus^19,20^, we compared the expression of the 75 CSPs in astrocytes with three major neuronal populations in the hippocampus: CA1 pyramidal neurons, CA2/CA3 pyramidal neurons and DG granule cells, using differential expression analysis. From each comparison, astrocyte-enriched and top three neuron-enriched CSPs were identified and ranked by log fold change (**Fig. 1D**). These comparisons identified 14 CSPs enriched in astrocytes, 10 of which were consistently enriched in two of the three comparisons: *Gpr37l1*, *Hepacam*, *Vcam1*, *Plxnb1*, *Ephb4*, *Slc3a2*, *Cadm4*, *Neo1*, *Lsamp* and *Alcam*. In addition, we found *Icam5*, *Epha6*, *Epha7*, *Shisa6* and *Nptx1* among the top three neuron-enriched CSPs.

Examination of their expression across the four cell type populations showed that *Gpr37l1* and *Hepacam* were expressed in a large proportion of astrocytes, exhibited high mean expression levels, and were highly specific to astrocytes (**Fig. 1E-F**). *Vcam1*, *Plxnb1 and Ephb4* were detected in a smaller fraction of astrocytes and displayed lower mean expression, yet they remained highly astrocyte-specific. *Slc3a2* and *Alcam* showed relatively uniform expression across the four cell types. *Cadm4* was expressed in astrocytes and CA1 neurons but exhibited low mean expression and was detected in a smaller fraction of cells. *Neo1* was expressed in astrocytes, CA1 neurons and DG granule cells. *Lsamp* exhibited high mean expression and was detected in a large fraction of both astrocytes and CA1 neurons.

Taken together, our hippocampal synaptic CSP proteome-informed scST analysis identifies 10 CSPs as candidates for astrocyte-expressed CSPs.

### Systematic profiling establishes GPR37L1, HepaCAM, and VCAM1 as high-confidence hippocampal astrocytic peri-synaptic CSPs

Next, we aimed to systematically profile the 10 candidate astrocyte-enriched CSPs, focusing on their spatial localization and expression patterns (**Fig. 2A-B**). We first investigated their spatial protein localization at low and high magnification in the hippocampus. To this end, we screened 21 antibodies targeting the 10 candidate astrocyte-enriched CSPs by IHC and western blot (**Table S1**). Antibodies exhibiting specific immunostaining patterns and detection of bands at the expected molecular weight were obtained for 8 candidate astrocyte-enriched CSPs (excluding PLXNB1 and EphB4). Of these, 7 – all except LSAMP – had previously been validated using knock-out or knock-down approaches (**Fig. 2B** and **Table S1**).

**Fig. 2.**
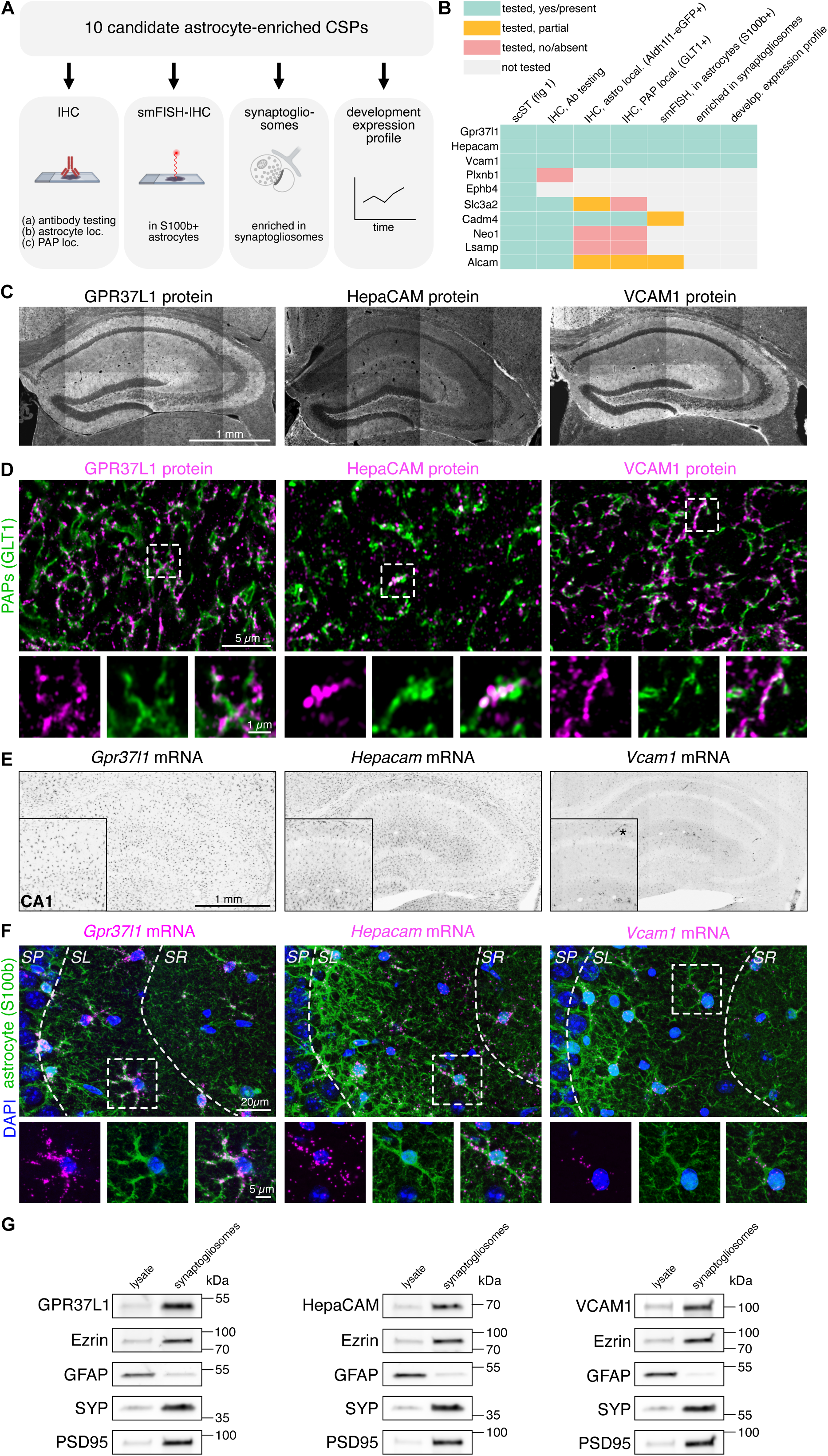
Systematic profiling of CSP candidates establishes GPR37L1, HepaCAM, and VCAM1 as high-confidence hippocampal astrocytic peri-synaptic CSPs. (A) Schematic of the workflow and experimental assays used to profile potential astrocytic peri-synaptic CSP candidates identified in Fig 1. Created with BioRender.com. (B) Summary heatmap showing candidate CSPs and outcomes across experimental assays used for profiling. (C) Low magnification single-plane immunostaining images of GPR37L1, HepaCAM and VCAM1 in the hippocampus of WT mice at P28. Representative of three independent experiments. (D) High magnification single-plane Airyscan super resolution images showing colocalization GPR37L1, HepaCAM and VCAM1 with the peri-synaptic astrocytic process marker GLT1 in the CA3 stratum lucidum (SL) of WT mice at P28. Insets highlight colocalization with GLT1. Representative of two independent experiments. (E) Low magnification single-plane smFISH images of *Gpr37l1*, *Hepacam* and *Vcam1* in the hippocampus of WT mice at P28. Insets highlight CA1 region. Representative of two independent experiments. Asterisk (*) in VCAM1 condition denotes presumable endothelial cell expression. (F) Representative smFISH z-stack images of *Gpr37l1*, *Hepacam* and *Vcam1* in the CA3 region of WT mice at P28. Insets highlight expression in astrocytes labeled with S100β within the CA3 SL. Hippocampal layers were delineated by VGLUT1 co-staining (not shown). Representative of two independent experiments. (G) Western blots of hippocampal lysates and hippocampal synaptogliosomes from P28 WT mice, probed for GPR37L1, HepaCAM and VCAM1 together with the peri-synaptic astrocytic process marker Ezrin, astrocytic cytosolic marker GFAP, presynaptic marker synaptophysin (SYP) and postsynaptic marker PSD95. Representative of two independent experiments. Molecular weight markers indicated in kDa.

Low magnification IHC revealed abundant expression of GPR37L1, HepaCAM, VCAM1, SLC3A2, CADM4 and NEO1 in the hippocampus, and not restricted to the hippocampal region (CA3) where we first identified these CSPs^18^ (**Fig. 2C** and **S2A**). Interestingly, LSAMP and ALCAM showed regional differences in expression: LSAMP was enriched in the CA1 region relative to the CA3 and DG, whereas ALCAM showed higher expression in CA1 and CA3 compared to DG (**Fig. S2A**). Using the Aldh1l1-eGFP astrocytic reporter mice, we found that GPR37L1, HepaCAM, VCAM1 and CADM4 showed strong localization to astrocytes, particularly in their fine processes (**Fig. S2B** and **S2C**). In contrast, SLC3A2 and ALCAM showed partial astrocytic localization, whereas NEO1 and LSAMP did not show detectable localization to Aldh1l1-eGFP labeled astrocytes (**Fig. S2C**).

To determine whether these astrocytic CSP candidates are localized to peri-synaptic astrocytic processes (PAPs), we performed co-immunostaining with the astrocytic glutamate transporter GLT1. Super-resolution Airyscan imaging revealed strong colocalization of GPR37L1, HepaCAM, VCAM1, CADM4 with PAPs (**Fig. 2D** and **S2D**). ALCAM showed partial colocalization to PAPs, whereas SLC3A2, NEO1 and LSAMP did not show colocalization (**Fig. S2D**).

Having shown that GPR37L1, HepaCAM, VCAM1, CADM4 and ALCAM localize to PAPs, we sought to further assess their transcript distribution in the hippocampus by using single-molecule fluorescent *in situ* hybridization (smFISH, RNAscope), a method with higher sensitivity than scST^21^. At low magnification, *Gpr37l1* and *Hepacam* exhibited a higher number of mRNA puncta compared to *Vcam1* (**Fig. 2E**), consistent with our spatial transcriptomics data (**Fig. 1E-F**). Based on spatial location, *Gpr37l1*, *Hepacam*, and *Vcam1* transcripts were largely absent from neuronal pyramidal layers (**Fig. 2E**), whereas *Cadm4* and *Alcam* transcripts were detected in pyramidal neurons and, in the case of *Alcam*, presumably also in inhibitory neurons (**Fig. S2E**). At higher magnification, co-staining with the astrocytic marker S100β confirmed localization of *Gpr37l1*, *Hepacam* and *Vcam1* transcripts within S100β+ astrocytic processes (**Fig. 2F**), while *Cadm4* and *Alcam* transcripts showed astrocytic and neuronal pyramidal layer localization (**Fig. S2F**).

We focused on GPR37L1, HepaCAM, and VCAM1 as astrocyte-enriched candidate CSPs at perisynaptic astrocytic processes (PAPs) and used an orthogonal approach to further confirm their localization within the tripartite synapse. To this end, we co-purified hippocampal PAPs together with hippocampal synaptosomes (synaptogliosome preparations^22,23^), and observed enrichment of GPR37L1, HepaCAM and VCAM1 in these fractions compared to total hippocampal lysate (**Fig. 2G**).

In the mouse hippocampus, synapse development progresses from initial synapse formation around P7, through a peak of synaptogenesis around P14, followed by a period of synapse refinement and pruning around P21, culminating in a mature state by P28 with well-established, functional neural circuits^24^. We therefore collected brains from wildtype littermates and examined the hippocampal protein expression of GPR37L1, HepaCAM and VCAM1 across postnatal developmental timepoints (P1-P7-P14-P21-P28) by western blotting (**Fig. S2G**). GPR37L1 expression was low at P1 and P7, drastically increased at P14 and remained high at P21 and P28. HepaCAM protein levels increased steadily throughout postnatal development, whereas VCAM1 expression peaked around P14.

In summary, we systemically profiled astrocytic CSP candidates using complementary approaches including IHC, smFISH-IHC, synaptogliosomes and developmental expression analyses. Together, these data identify GPR37L1, HepaCAM, and VCAM1 as astrocyte-enriched CSPs across the hippocampus and localized at peri-synaptic astrocytic compartments, positioning them to regulate synapse development and function.

### Synaptic VCAM1 and GPR37L1 complexes contain neuronal and astrocytic CSPs

Our candidate profiling revealed astrocytic perisynaptic localization of GPR37L1, HepaCAM and VCAM1, suggesting that these proteins may participate in interactions at the tripartite synapse. Because astrocytic HepaCAM has previously been shown to localize to synapses and regulate inhibitory synapse development^25^, we focused subsequent analyses on GPR37L1 and VCAM1.

To define the molecular context in which these proteins operate at the tripartite synapse, we next mapped their synaptic interactomes through co-immunoprecipitation (co-IP) of endogenous VCAM1 or GPR37L1. We first prepared hippocampal crude synaptosomes from P21 mouse brains and then performed co-IP for VCAM1 or GPR37L1 (**Fig. 3A** and **S3A**). The subsequently eluted proteins were analyzed by liquid chromatography-tandem mass spectrometry (LC-MS/MS) using data-independent acquisition (DIA) and quantified using a label-free approach (**Fig. 3A** and **S3B**). For each biological replicate, the crude synaptosomal sample was split into two equal fractions for IP and a species-matched IgG IP control condition.

**Fig. 3.**
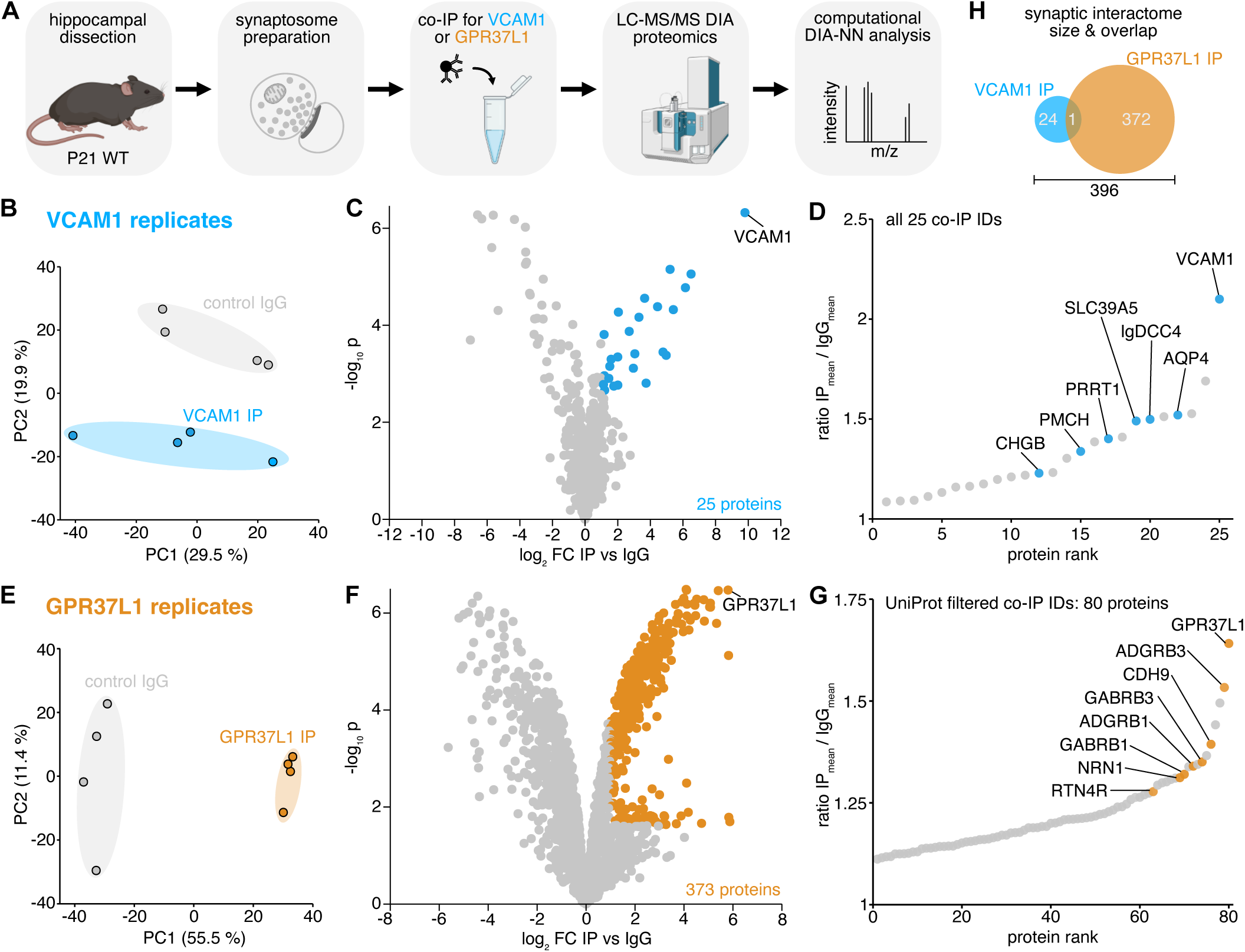
Proteomics reveals distinct hippocampal synaptic interactomes of VCAM1 and GPR37L1. (A) Schematic of the workflow used to dissect the hippocampal synaptic interactome of GPR37L1 and VCAM1. Created with BioRender.com. (B) PCA of control IgG and VCAM1 IP samples showing clear separation by condition (n = 4 biological replicates). (C) Volcano plot showing differentially expressed proteins (DEPs) between VCAM1 IP and control IgG identified by Limma (adj. p < 0.05; log_2_FC > 1). (D) Co-immunoprecipitated (co-IP) proteins ranked by IP (mean) / IgG (mean) ratio. (E) PCA of control IgG and GPR37L1 IP samples showing clear separation by condition (n = 4 biological replicates). (F) Volcano plot showing differentially expressed proteins (DEPs) between GPR37L1 IP and control IgG identified by Limma (adj. p < 0.05; log_2_FC > 1). (G) Co-immunoprecipitated (co-IP) proteins filtered based on UniProt cell surface proteins and ranked by IP (mean) / IgG (mean) ratio, same highlighted proteins as in panel (F).

Protein identification and quantification performed with DIA-NN software yielded more than 1700 different proteins across VCAM1 co-IP and IgG samples after filtering (**Fig. S3B** and **S3E-S3F**). PCA analysis showed clear separation and clustering by condition (**Fig. 3B**). VCAM1 was exclusively detected in VCAM1 co-IP samples, and not in IgG controls, confirming the specificity of the pull-down (**Fig. S3C** and **S3D**).

Differential expression analysis identified 25 proteins significantly enriched in the VCAM1 co-IP condition, with VCAM1 itself showing the strongest enrichment (**Fig. 3C and 3D**). As we were especially interested in protein-protein interactions with potential roles in cell-cell communication at the synapse, we limited our subsequent analysis to CSPs based on UniProt annotations. To identify potential VCAM1 interactors, we therefore filtered the differentially expressed proteins for their cell surface localization using UniProt annotations (**Fig. S3B** and **3C-D)**. From this list, IgDCC4 and SLC39A5 have no described role in the brain. Interestingly, we identified two neuropeptides, CHGB and PMCH, in the VCAM1 co-IP.

Comparisons with a recently published synaptic proteome study^26^ revealed that over 40% of our identified interactors were previously reported in the hippocampal synaptic proteome (**Fig. S3G**). Further characterization using the synaptic protein database described by Sorokina, Mclean, Croning et al.^27^, which compiles synaptic proteins identified in proteomic studies from the past 20 years and integrates multiple external resources, showed that more than 70% of the enriched proteins were previously found in synaptosomes (**Fig. S3G**). We cross-referenced these proteins with a curated synaptic protein database SynGO^28^ and found one neuronal membrane protein: PRRT1 (**Fig. S3G).** PRRT1 is an AMPA receptor-associated protein that is required for hippocampal excitatory synapse development and function^29^.

For GPR37L1 IP we identified more than 2100 different proteins across co-IP and IgG samples after filtering, with consistently more proteins identified in the co-IP samples (**Fig. S3B** and **S3I-J**). PCA analysis showed clear separation of GPR37L1 co-IP samples from IgG controls and tight clustering by condition (**Fig. 3E**). Consistent with the specificity of the pull-down, GPR37L1 itself was detected exclusively in the GPR37L1 co-IP samples and not in IgG controls (**Fig. S3H**).

Differential expression analysis identified 373 proteins significantly enriched in the GPR37L1 co-IP condition, substantially more than the number of proteins detected for VCAM1 (**Fig. 3F and 3G**). Comparison with a recently published hippocampal synaptic proteome study^26^ revealed that over 50% of the coIP proteins had previously been reported to localize at synapses (**Fig. S3K**). Futhermore, cross-referencing with the synaptic protein database compiled by Sorokina, Mclean, Croning et al.^27^ showed that more than 90% of the GPR37L1 coIP had previously been identified in synaptosomes (**Fig. S3K**).

Consistent with this synaptic enrichment, SynGO analysis indicated that approximately 20% of the GPR37L1 co-IP proteins have annotated synaptic functions (**Fig. S3K** and **S3L**). Indeed, gene ontology analysis further highlighted Biological Process (BP) terms associated with synaptic organization, including “presynapse organization”, “adherens junction organization”, and “calcium-dependent cell-cell adhesion via plasma membrane cell adhesion molecules” (**Fig. S3M**).

To focus on potential synaptic signaling components, we next filtered the GPR37L1 co-IP proteins based on their cell surface localization using UniProt annotations, identifying a repertoire of 80 CSPs (**Fig. 3G** and **S3B**). We ranked these CSPs according to their IP/IgG ratio and highlighted the proteins in the top 15 with a known synaptic functional role. Notably, we found two adhesion GPCRs with well known roles in synaptic development: ADGRB3^30,31^ and ADGRB1^32,33^. Interestingly, we also identified RTN4R, a well known receptor^34^ through which ADGRB3 enables hippocampal excitatory synapse formation^35^. Neuronal CDH9 was previously shown to regulate hippocampal synapse development in the CA3 region^36^, which is the same region as our source proteomic FASS dataset. Other top hits included GABRB3 and GABRB1, which regulate inhibitory synaptic tranmission^37,38^, and NRN1, an activity regulated protein involved in dendritic spine development and stabilization in the hippocampal DG^39,40^.

Comparison of the GPR37L1 and VCAM1 interactomes revealed minimal overlap, with only a single shared intracellular protein (SSBP1). Together, these analyses indicate that GPR37L1 and VCAM1 occupy distinct molecular environments at hippocampal synapses.

### Multi-metric image-based analysis pipeline identifies VCAM1 as a regulator of hippocampal synaptic structural parameters

Recent findings indicate that GPR37L1 modulates astrocyte morphological complexity in the cortex^41^, and synaptic transmission in the spinal cord dorsal horn^42^. Together with our identification of the hippocampal synaptic interactome for GPR37L1, these findings suggest that astrocytic GPR37L1 may contribute to synaptic development in the hippocampus. To our knowledge, the function of astrocytic VCAM1 in synaptic development has not been explored.

To investigate GPR37L1 and VCAM1’s role in hippocampal synaptic development, we used a adeno-associated viral-vector (AAV)-based CRISPR/Cas9-mediated knockout (KO) strategy coupled with a custom-developed synapse image analysis pipeline. Our experimental approach involves an injection of AAV (AAV2/5-U6::gRNA-CAG::FLEX-Lck-GFP) containing guide RNAs (gRNAs) targeting *Gpr37l1* or *Vcam1* in one hemisphere and a control AAV harboring a LacZ gRNA in the contralateral hemisphere of constitutive Cas9 expressing mice (H11-Cas9)^43^ (**Fig. 4A**).

**Fig. 4.**
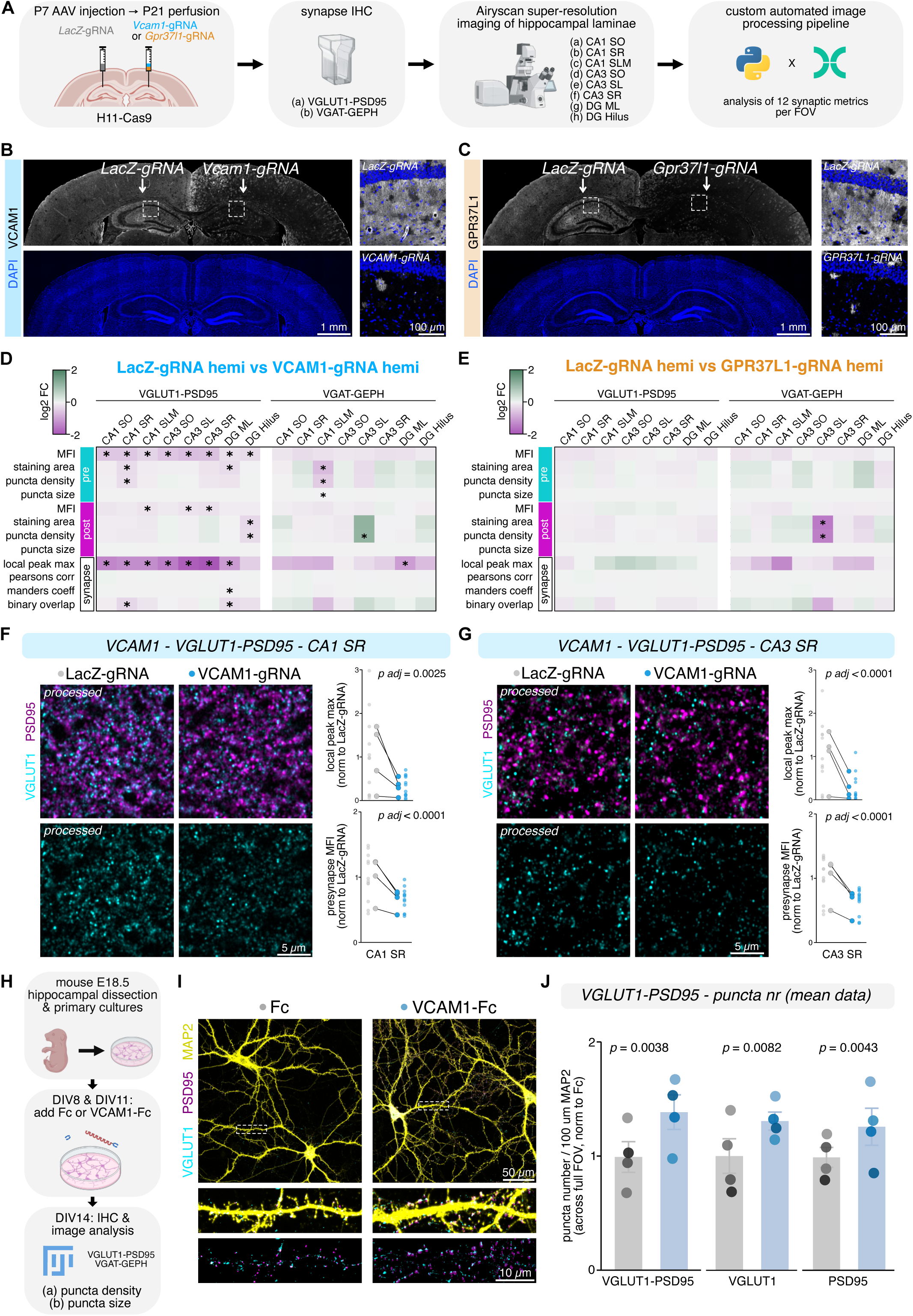
Multi-metric lamina-resolved image analysis identifies VCAM1, but not GPR37L1, as a regulator of hippocampal synapse development. (A) Schematic of the workflow used to assess the effect of constitutive CRISPR/Cas9-mediated KO of *Vcam1* or *Gpr37l1* on hippocampal synapse development. SO: stratum oriens, SR: stratum radiatum, SLM: stratum lacunosum-moleculare, SL: stratum lucidum, DG: dentate gyrus, ML: molecular layer. Created with BioRender.com. (B) Quality control of *Vcam1* KO by immunohistochemistry for VCAM1 and DAPI. Insets show CA1 region of both hemispheres. (C) Quality control of *Gpr37l1* KO by immunohistochemistry for GPR37L1 and DAPI. Insets show CA1 region of both hemispheres. (D) Heatmap showing the log2 fold change in synaptic metrics comparing LacZ-gRNA and VCAM1-gRNA hemispheres across hippocampal laminae. For each synaptic metric and lamina, 3 FOVs (sections) per brain from 4 brains were analyzed. Log2-transformed metrics were analyzed using a linear mixed-effects model with gRNA, hippocampal lamina and their interaction as fixed effects to test gene- and lamina-specific effects, and brain as a random intercept to account for inter-brain variability. Asterisks denote significant differences after false discovery rate (FDR) correction for each metric across hippocampal laminae (FDR < 0.05). (E) Heatmap showing the log2 fold change in synaptic metrics comparing LacZ-gRNA and GPR37L1-gRNA hemispheres. Same sampling, plotting and analyses as in (G). (F) Representative processed Airyscan images of VGLUT1 and PSD95 in CA1 SR from LacZ-gRNA- and VCAM1-gRNA-injected hemispheres, with quantification of local peak max and presynapse MFI normalized to the LacZ-gRNA hemisphere. Each small data point represents a single FOV; larger linked data points represent hemisphere means. Statistical significance was assessed using a linear mixed effects model as in (G), followed by Tukey’s post hoc tests and FDR corrected significance. (G) Representative processed Airyscan images of VGLUT1 and PSD95 in CA3 SR from LacZ-gRNA- and VCAM1-gRNA-injected hemispheres, with quantification of local peak max and presynapse MFI normalized to the LacZ-gRNA hemisphere. Same sampling, plotting and analyses as in (F). (H) Schematic of the workflow used to assess the effect of VCAM1-Fc protein treatment on hippocampal synapses *in vitro*. Created with BioRender.com. (I) Representative images of hippocampal neurons treated with recombinant protein from DIV8 to DIV14. Fc or VCAM1-Fc (2 µg/mL) was added at DIV8 and replenished at DIV11. Neurons were fixed at DIV14 and stained for VGLUT1 (cyan), PSD95 (magenta), and MAP2 (yellow). Insets show the merge followed by the isolated VGLUT1-PSD95 signal. (J) Quantification of excitatory synaptic puncta density along MAP2+ dendrites. VGLUT1+PSD95+ colocalized puncta, as well as VGLUT1+ and PSD95+ puncta separately, were measured per 100 μm dendrite and normalized to the Fc condition. Each FOV contained two neuronal somas for standardized sampling; 9-11 FOVs per condition were analyzed in N = 4 independent experiments. Each data point represents one experiment is color-shaded by experiment. A linear mixed-effects model on log2 transformed data was used to account for inter-experimental variability. Post-hoc pairwise comparisons were conducted using Tukey’s HSD test. Individual FOV data points are shown in Fig. S5B.

To assess the KO efficiency of CRISPR gRNAs targeting *Gpr37l1* and *Vcam1*, we first used cultured hippocampal astrocytes. Western blot analysis of WT astrocyte lysates confirmed the expression of VCAM1 in primary astrocytes (**Fig. S4A**). In contrast, GPR37L1 showed minimal expression in cultured astrocytes. We therefore restricted *in vitro* testing to gRNAs targeting *Vcam1*. We infected astrocyte cultures from H11-Cas9 mice with AAVs harboring GFP (transduction control), a gRNA targeting *LacZ* (gRNA#1 or #2), or a gRNA targeting *Vcam1* (gRNAs #1-3) at 14 days in vitro (DIV14) (**Fig. S4B**). After two weeks, lysates were collected and analyzed by western blot. All three *Vcam1* gRNAs effectively reduced VCAM1 protein levels compared to cells infected with the LacZ control gRNAs.

Because GPR37L1 expression was too low in cultured astrocytes to reliably assess gRNA efficiency, we evaluated the activity of a single Gpr37l1 gRNA directly in vivo by immunohistochemistry (IHC) using the strategy described above. Using this approach, we observed a clear qualitative reduction of both endogenous VCAM1 (gRNA #2) and GPR37L1 protein levels in the ipsilateral hippocampus compared to the respective contralateral LacZ-injected control (**Fig. 4B-C**). This depletion was confirmed for every section and field-of-view (FOV) included in downstream analyses as a quality control measure. Notably, sparse astrocyte territories retaining detectable VCAM1 or GPR37L1 immunoreactivity were still observed (**Fig. 4B-C**).

Having established a clear CRISPR/Cas9-mediated reduction of VCAM1 and GPR37L1 protein levels *in vivo*, we proceeded with our *in vivo* loss-of-function (LOF) approach to assess the effect of these CSPs on hippocampal synapses hippocampal synapses. AAVs harboring gRNAs were injected at P7, a developmental timepoint at the onset of synaptogenesis and late enough to not impact the astrocyte progenitors derived from radial glial cells^44^ (**Fig. 4A**). At P21, mice were perfused, brains were collected, coronally sectioned and immunostained for excitatory pre- and postsynaptic markers (VGLUT1+; PSD95+) and inhibitory synaptic markers (VGAT+; GEPH+) (**Fig. 4A**). The P7-P21 interval encompasses the expression peak period of both VCAM1 and GPR37L1 hippocampal protein expression as shown in **Fig. S2G**.

As both VCAM1 and GPR37L1 show broad expression throughout the hippocampus, high magnification Airyscan super-resolution images were acquired from the ipsi- and contralateral hippocampus for 8 synaptic regions: CA1 *Stratum Oriens* (SO), CA1 *Stratum Radiatum* (SR), CA1 *Stratum Lacunosum Moleculare* (SLM), CA3 SO, CA3 *Stratum Lucidum* (SL), CA3 SR, DG *Molecular Layer* (ML) and DG *Hilus* (**Fig. 4A**). We developed a custom high-throughput computational synapse analysis pipeline to measure 12 commonly analyzed synaptic metrics: mean fluorescence intensity (MFI), staining area, puncta density and puncta size for both pre-and postsynaptic images per field of view (FOV) (**Fig. S4C**). For colocalization assessment, the pipeline measures Pearsons’s correlation coefficient, Manders’ overlap coefficient, overlap between binarized pre- and postsynapse images, and performs a custom analysis evaluating the colocalization of pre- and postsynaptic local intensity peak maxima (**Fig. S4C** and **S4D**).

The pipeline generated high-dimensional, multi-metric synaptic datasets for VCAM1 and GPR37L1. Linear mixed-effects modeling was subsequently applied, and dimensionality reduction via PCA demonstrated a clear separation and clustering of VCAM1-gRNA- and LacZ-gRNA-injected hemispheres based on synaptic metrics (**Fig. S5A** and **S5B**). In contrast, the GPR37L1 dataset showed less distinct separation and clustering between hemispheres (**Fig. S5C** and **S5D**).

To determine whether synaptic metrics differed significantly between VCAM1-gRNA- or GPR37L1-gRNA- and LacZ-gRNA-injected hemispheres, we performed pairwise comparisons for each synaptic metric. In the VCAM1 excitatory synapse (VGLUT1-PSD95) dataset, *Vcam1* KO resulted in a consistent downward trend across nearly all synaptic metrics (**Fig. 4D** and **S5B**). At least 2 synaptic metrics were significantly reduced in every hippocampal lamina in the VCAM1-gRNA-injected hemisphere compared to the LacZ-gRNA injected control hemisphere. As an illustrative example, in CA1 SR, analysis of local peak maxima revealed a reduction in apposed pre- and postsynaptic puncta as well as reduced VGLUT1 MFI (**Fig. 4F**). In another example with CA3 SR, we also observed a significant reduction in local peak maxima in apposed pre- and postsynaptic puncta and VGLUT1 MFI. However, not all synaptic metrics differed significantly, as illustrated by PSD95 MFI in CA3 SO (**Fig. S6A**).

For the VCAM1 inhibitory synapse (VGAT-GEPH) dataset, we did not observe a uniform trend across all synaptic metrics (**Fig. 4D** and **S5B**). However, CA1 SLM stood out, with 3 synaptic metrics significantly altered, namely VGAT staining area, puncta density and puncta size (**Fig. 4D** and **S6B**). An interesting observation was the increased GEPH puncta density in CA3 SL (**Fig. 4D**). However, closer inspection of the data suggests that this effect may be driven by highly variable data points across individual brains. (**Fig. S6C**).

For the GPR37L1 excitatory (VGLUT1-PSD95) and inhibitory (VGAT-GEPH) synapse dataset, we did not observe a uniform trend across synaptic metrics (**Fig. 4E**), as illustrated by VGLUT1-PSD95 in CA1 SLM with Pearson’s correlation coefficient (**Fig. S6D**). However, GEPH staining area and puncta density in the CA3 SL were significantly reduced following GPR37L1 depletion (**Fig. 4E** and **Fig. S6E**).

Taken together, these findings demonstrate that the loss of VCAM1 impairs excitatory synaptic development in the hippocampus and, to a lesser extent, also affects inhibitory synaptic development in specific hippocampal laminae. In contrast, loss of GPR37L1 had a minimal effect on hippocampal synaptic development.

### Recombinant VCAM1 increases excitatory but not inhibitory hippocampal synapse density in primary hippocampal cultures

To complement our *in vivo* findings following CRISPR/Cas9-mediated *Vcam1* KO, we employed a simplified and independent *in vitro* assay. To this end, we used cultured hippocampal neurons to test whether addition of the recombinant, soluble VCAM1 ectodomain affects synapse formation (**Fig. 4H**).

Due to poor neuronal health in the absence of astrocytes, we co-cultured neurons with a glial feeding layer in a “sandwich” configuration, preventing direct contact between astrocytes and neurons. Recombinant VCAM1-Fc or Fc control protein, which were verified by western blot, was added to the culture medium at days *in vitro* (DIV) 8 and 11 to assess synaptogenic effects (**Fig. S7A**). Neurons were fixed at DIV14 and immunostained for either excitatory synaptic markers (VGLUT1 and PSD95) or inhibitory synaptic markers (VGAT and GEPH) and the dendritic marker MAP2. Confocal images were acquired and puncta density and size per field of view (FOV) were quantified for pre-, post, and colocalized synaptic markers using a semi-automatic analysis pipeline (distinct from the pipeline used in Fig. 3) (**Fig. 4H**).

Treatment with VCAM1-Fc significantly increased the density of VGLUT1+, PSD95+, and colocalized VGLUT1+/PSD95+ puncta compared to the Fc control condition (**Fig. 4I-J** and **S7D**). No differences in pre- or postsynaptic puncta size were detected between VCAM1-Fc and Fc-treated cultures (**Fig. S7F**). In contrast, analysis of inhibitory synapses (VGAT-GEPH) revealed no significant differences in puncta density or size between conditions (**Fig. S7B-C, S7E** and **S7G**).

Taken together, these results demonstrate that addition of recombinant, soluble VCAM1 ectodomain is sufficient to support excitatory, but not inhibitory, hippocampal synapse formation *in vitro*.

## DISCUSSION

Here, through a systematic cross-modal approach, we identified VCAM1 as an astrocyte-expressed protein that localizes to PAPs and regulates hippocampal excitatory synapse development both *in vivo* and *in vitro*.

We leveraged a synapse proteomics resource previously generated in our lab^18^ in an integrative, systems-level framework. By combining targeted single-cell spatial transcriptomics with protein-level validation, we distilled a large synaptic CSP list into a set of high-confidence astrocytic peri-synaptic proteins. This stepwise prioritization was critical, as transcript detection alone does not necessarily predict protein localization or abundance. Indeed, several candidate CSPs showed discordance between mRNA and protein levels or lacked peri-synaptic localization, underscoring the importance of our multi-modal approach. For example, *Vcam1* transcripts were qualitatively relatively low despite robust protein detection (noting antibody dependence), whereas *Hepacam* transcripts were comparatively abundant but showed lower protein levels.

Our data establish GPR37L1, HepaCAM and VCAM1 as bona fide astrocytic peri-synaptic CSPs in the hippocampus. We prioritized two candidates for further investigation; VCAM1 and GPR37L1, as HepaCAM was previously found to be implicated in synaptic development. While VCAM1 is best known for its role in immune cell adhesion and endothelial cell biology^45^, our findings reveal a previously unrecognized role at hippocampal tripartite synapses. In both our scST and smFISH analyses, *Vcam1* expression in astrocytes appeared qualitatively low. In the scST dataset, *Vcam1* transcripts were detected in only a subset of astrocytes, whereas the more sensitive smFISH approach revealed robust *Vcam1* transcripts across astrocytes. smFISH combined with IHC further localized VCAM1 transcripts to S100β+ astrocytic processes. At the protein level, VCAM1 was detected in association with the astrocytic glutamate transporter GLT1 at PAPs and was also present in synaptogliosome preparations. Moreover, the synaptic VCAM1 interactome suggests a potential role in synapse regulation. Together, these observations suggest that VCAM1 is positioned within astrocytic PAPs to influence synaptic organization.

The *in vivo* CRISPR/Cas9-mediated loss-of-function (LOF) experiments demonstrate that VCAM1 is required for proper excitatory synapse development across multiple hippocampal synaptic regions. Our multi-metric, lamina-resolved analysis pipeline proved particularly powerful in this context. Rather than relying on a single synaptic readout, we quantified commonly analyzed metrics of synaptic structure, including puncta density, intensity, size, and multiple colocalization metrics. The convergence of these measures revealed a consistent downward shift in excitatory synaptic metrics upon VCAM1 deletion, strengthening the robustness of the phenotype. In contrast, inhibitory synapses exhibited more modest and lamina-restricted changes, with the most pronounced effects in CA1 SLM. The preferential effect on excitatory synaptic development was independently supported by our *in vitro* gain-of-function assay, in which the recombinant VCAM1 ectodomain selectively enhanced excitatory, but not inhibitory, synapse density. Combined, these observations suggest that VCAM1 primarily regulates excitatory synapse development.

In contrast to *VCAM1*, *Gpr37l1* expression in astrocytes appeared markedly higher in both our scST and smFISH analyses, approaching levels typical of astrocytic marker genes. In the scST dataset, *GPR37L1* transcripts were detected at high levels in astrocytes, a finding that was confirmed by the smFISH approach. smFISH combined with immunohistochemistry further localized abundant *Gpr37l1* transcripts to S100β+ astrocytic processes. At the protein level, GPR37L1 was detected in association with the astrocytic glutamate transporter GLT1 at PAPs and was also present in synaptogliosome preparations. In addition, the unexpectedly extensive synaptic GPR37L1 interactome points to a potential role in synapse regulation. However, in the *in vivo* CRISPR/Cas9-mediated LOF experiments we only found a lamina-restricted inhibitory postsynaptic developmental phenotype in the CA3 SL, and we did not observe a consistent trend across all synaptic metrics for inhibitory synapses.

Interestingly, astrocytic GPR37L1 was recently shown to regulate excitatory and inhibitory synaptic transmission in the spinal dorsal horn^42^ and modulate astrocyte morphological complexity in the cortex^41^. Thus, together with our extensive synaptic interactome, GPR37L1 may have a role in synaptic function rather then synaptic development. Future studies using conditional, astrocyte-specific deletion combined with electrophysiological approaches will be necessary to determine whether GPR37L1 modulates hippocampal synaptic transmission.

Our proteomic analysis of the VCAM1 synaptic interactome provides insight into its potential function at synapses. Among the potential VCAM1 interacting proteins was PRRT1, which has been shown to regulate AMPA receptor trafficking and hippocampal excitatory synapse function underlying higher-order cognitive plasticity^29^. Notably, PRRT1 has been proposed to maintain a pool of extrasynaptic AMPARs necessary for synaptic transmission and plasticity. However, it remains unclear whether VCAM1 directly binds PRRT1 or instead associates with it in a larger multi-protein complex. Another intriguing candidate interactor of VCAM1 is the neuropeptide PMCH, encoding the prepro-melanin-concentrating hormone (MCH). Interestingly, PMCH is not expressed by hippocampal neurons but by hypothalamic neurons that project MCH-rich terminals to the hippocampal CA1 area^46^. We previously showed that treatment of hippocampal neurons with MCH peptide strongly reduces synaptic transmission and synaptic strength^47^. One possibility is that astrocytic VCAM1 locally sequesters MCH peptides; disruption of this interaction in the VCAM1 KO could therefore contribute to the observed developmental reduction in synapse number. We also identified the neuropeptide-associated protein CHGB, a luminal component of neuronal dense-core vesicles that is secreted into the extracellular space during activity-dependent vesicle release. Similarly, VCAM1 could sequester CHGB peptides, although double (CHGB and CHGA) KO does not affect synapse density in cultured hippocampal neurons, thus likely not explaining our VCAM1 KO synaptic phenotype.

An important question is whether VCAM1 acts primarily as a membrane-attached protein at astrocytic processes or through a soluble, cleaved form. In our *in vitro* recombinant protein experiments, neurons were cultured on an astrocytic feeder layer, making it difficult to determine whether VCAM1 acts directly at excitatory synapses or indirectly through astrocyte-mediated mechanisms, such as the release of synaptogenic factors. Notably, VCAM1 can undergo ectodomain shedding in the brain^45^, raising the possibility that a soluble form of VCAM1 may contribute to its synaptic effects. Together, these observations suggest that both contact-dependent and secreted VCAM1 signaling could modulate synaptic development.

Analysis of the GPR37L1 hippocampal synaptic interactome revealed a surprisingly large number of potential interactors (373 proteins), including components of protein complexes. The pull-down was highly specific as we used a GPR37L1 KO-validated antibody. In contrast, the VCAM1 pull-down, also performed with a KO-validated antibody, identified only 25 proteins. One possible explanation for this difference is the clonality of the antibodies used: a monoclonal antibody was used for GPR37L1, whereas a polyclonal antibody was used for VCAM1, which may lead to epitope masking and reduced recovery of interacting proteins.

In the hippocampal synaptic interactome of GPR37L1, one of the most prominent complexes found was the ADGRB-RTN4R complex^34^. ADGRB1 and ADGRB3, but not ADGRB2, have been show to bind RTN4R *in trans* to regulate synapse formation^35^. Strikingly, our GPR37L1 coIP dataset contained ADGRB1 and ADGRB3 but not ADGRB2. However, ADGRB1 and ADGRB3 are known to regulate excitatory synapse formation and synaptic transmission^35^, which does not align with the inhibitory postsynaptic developmental phenotype observed in CA3 SL. In contrast, we identified GABRB3 and GABRB1 in our GPR37L1 coIP dataset, both of which regulate inhibitory synaptic tranmission^37,38^, and are therefore more consistent with our phenotype. Another notable observation was that GPR37L1 co-immunoprecipited multiple members of the cadherin family, which regulate neural circuit formation^48^. These included CDH4, CDH6, CDH8, CDH9, CDH10 and CDH11. One of the top GPR37L1 coIP hits was CDH9, which has previously been implicated in excitatory synaptic development in the CA3 SL^36^ and CA1 SR^49^. Additionally, CDH6 and CDH10 are required for excitatory synaptic development and plasticity in the CA1 SO and SR^49^. CDH11 has also been reported to regulate GABAergic inhibitory synapse development in cultured hippocampal neurons^50^.

Our findings expand the repertoire of astrocytic CSPs implicated in hippocampal synaptic development beyond the previously described roles of Ephrin-B1^15,16^ and CD38^17^. By leveraging our lamina-resolved synapse analysis, we can directly compare layer-specific phenotypes to those identified in Ephrin-B1 and CD38 KO models. Astrocytic Ephrin-B1 conditional knockout (cKO) increased VGLUT1+ puncta and VGLUT1+ PSD95+ excitatory synapses in the CA1 SR, whereas VCAM1 KO produced the opposite effect, decreasing VGLUT1+PSD95+ synapses in CA1 SR, primarily through a reduction in VGLUT1+ presynaptic metrics. Astrocytic CD38 cKO showed a reduction in excitatory VGLUT1+PSD95+ synaptic density in the CA1 SR, similar to VCAM1 KO. Whether this effect in the CD38 cKO is mainly driven by VGLUT1 or PSD95 is not known, however. These findings suggest that distinct astrocytic CSPs can exert divergent regulatory effects on excitatory synapse density within the same hippocampal lamina. Notably, Ephrin-B1 deletion did not affect GAD65+ inhibitory synapses in CA1 SR or SLM. In contrast, VCAM1 deletion reduced VGAT+GEPH+ inhibitory synapses in CA1 SR and produced an even more pronounced reduction in CA1 SLM synaptic metrics, largely driven by decreased VGAT+ presynaptic staining. Together, our findings indicate that astrocytic CSPs differentially regulate excitatory and inhibitory synapse development, potentially in a lamina-specific manner in interaction with neuronal lamina-specific CSPs^51^.

Future work combining astrocyte-specific manipulations with electrophysiological recordings will be essential to determine how VCAM1 affects synaptic transmission, and plasticity. More broadly, our cross-modal strategy; integrating targeted spatial transcriptomics, protein-level validation, *in vivo* CRISPR/Cas9 perturbation, high-content image processing and analysis, *in vitro* functional assays, and synaptic proteomic interactomics, provides a generalizable framework to uncover glial regulators of circuit assembly. In conclusion, we identify astrocytic VCAM1 as a regulator of hippocampal excitatory synapse development.

### Limitations of the study

Our astrocytic CSP profiling relied on the glutamate transporter GLT1 as a marker for peri-synaptic astrocytic processes (PAPs), which biased the analysis toward excitatory synapses. Because astrocytes also express GABA transporters such as GAT1 and GAT3, astrocytic CSPs associated with inhibitory synapses are likely underrepresented. For *in vivo* LOF studies, we used a strategy to constitutively knockout VCAM1 and GPR37L1. Although astrocytic PAP localization was extensively validated, we cannot exclude contributions from other cell types expressing these proteins. While VCAM1 and GPR37L1 KO efficiency was robustly validated, we did not include a marker to identify AAV-transduced astrocytes. This limitation was particularly relevant for the LacZ control condition, as we could not verify whether the FOVs corresponds to astrocytes that were successfully transduced with the LacZ control virus.

## RESOURCE AVAILABILITY

### Lead contact

Requests for further information and resources should be directed to and will be fulfilled by the lead contact, Joris de Wit.

### Material availability

This study did not generate new unique reagents.

### Data availability

- The raw and intermediate targeted imaging-based single-cell spatial transcriptomics data have been deposited to the BioImage Archive: S-BIAD2879.
- The raw and intermediate mass spectrometry proteomics data have been deposited to the ProteomeXchange Consortium via the PRIDE database: PXD070371 (VCAM1 IP) and PXD PXD070422 (GPR37L1 IP).
- The raw and intermediate CRISPR/Cas9 KO *in vivo* synapse imaging data have been deposited to the BioImage Archive: S-BIAD2530.
- The raw and intermediate *in vitro* synapse imaging data have been deposited to the BioImage Archive: S-BIAD2430.

### Code availability

To ensure full reproducibility and transparency, all processing pipelines, analysis code and code dependencies for scST, MS, *in vivo* synapse imaging data and *in vitro* synapse imaging data are publicly available at https://github.com/cgeyskens/phd, accompanied with experiment-specific documentation.

## AUTHOR CONTRIBUTIONS

C.G., D.D. and J.d.W conceived the project and designed the experiments. C.G. prepared samples for scST experiments and analyzed scST data with guidance from F.P. C.G. performed experiments to profile astrocyte-enriched CSP candidates. C.G. and K.B. prepared crude synaptosomes for MS experiments. L.L. and P.M. prepared samples and performed MS experiments. C.G. analyzed the MS data with guidance from P.M. and D.D. E.L. produced AAVs and performed AAV stereotactic injections for *in vivo* synapse analysis. C.G. performed the *in vivo* synapse analysis experiments, developed the computational synapse image analysis pipeline and conducted downstream analyses. C.G. and E.K. performed *in vitro* synapse quantification experiments and analyzed the data. J.V. maintained mouse colonies and performed genotyping. C.G., D.D. and J.d.W wrote the manuscript with input from all authors.

## ACKNOWLEDGEMENTS

We thank Marinka Brouwer, Stephanie Chanda and Beatriz Marques for help with primary neuronal cultures. We thank Steffen Fieuws (Interuniversity Institute for Biostatistics and Statistical Bioinformatics, Leuven, Belgium) for advice on statistical modeling. We thank Benjamin Pavie (VIB-Bioimaging Core, Center for Neuroscience) for input on the synapse image analysis processing pipeline. We thank the VIB-Bioimaging Core, Center for Neuroscience for microscopy training and advice. The resources and services used in this work were provided by the VSC (Flemish Supercomputer Center), funded by the Research Foundation - Flanders (FWO) and the Flemish Government. D.D. is supported by Fonds Wetenschappelijk Onderzoek (FWO) Postdoctoral fellowship 12W5218N, 12W5221N and FWO Research Grant 1513320N; J.d.W. is supported by FWO Project Grants G0C4518N, G0A8720N, G0A8320N; FWO EOS Grant G0H2818N, and ERANET-NEURON TAO2PATHY Grant G0I3118N. Some figure panels were made in BioRender, as indicated in corresponding figure legends.

## MATERIALS AND METHODS

### Animals

All animal experiments were conducted according to KU Leuven guidelines and approved by an ethical commission (ECD project: P191/2021). Mice were maintained in a specific pathogen-free facility under standard housing conditions with continuous access to food and water. The health and welfare of the animals was supervised by a designated veterinarian. Wild-type (WT) C57BL/6J mice originated from Charles River. Aldh1l1-eGFP mouse line was obtained from Dr. Matthew Holt (VIB CBD, Leuven). H11-Cas9 mouse line was obtained from Dr. Jan Cools (VIB CCB, Leuven). Genotypes were regularly checked by polymerase chain reaction (PCR) analysis.

### Plasmids

An AAV backbone vector was generated using pAAV-U6-sgRNA(SapI)_hSyn-GFP-KASH-bGH (SpGuide acceptor; Addgene #60958) and AAV pCAG-FLEX-tdTomato-WPRE (Addgene #51503). The FLEX-tdTomato-WPRE cassette was excised and subcloned into pAAV-U6-sgRNA(SapI) using XhoI and XbaI restriction enzymes, yielding pAAV-U6::gRNA-CAG::FLEX-Lck-GFP. Guide RNA (gRNA) sequences targeting *Vcam1* and *Gpr37l1* for gene KO were designed using CRISPick database, with the Mouse GRCm38 (NCBI RefSeq) chosen as reference genome, CRISPRko as mechanism and the SpyoCas9 (Hsu 2013 et al.^52^) tracrRNA as enzyme. For each target gene, the top three ranked gRNA sequences were chosen. The LacZ gRNA sequences were obtained from Shuster et al. 2022^53^ (**Table S5**). Forward and reverse oligos were designed and SapI (NEB) restriction site overhangs were added. gRNA sequences were cloned into pAAV-U6::gRNA-CAG::FLEX-GFP using SapI restriction enzyme. AAVs were produced using pRep/Cap (AAV2/5, Addgene #232922) and pΔF6 (Addgene #112867) helper plasmids using previously established iodixanol purification.

### Cell lines

HEK293T cells were obtained from American Type Culture Collection (cat# CRL-11268) and grown in Dulbecco’s modified Eagle’s medium (DMEM, Invitrogen) supplemented with 10% fetal bovine serum (FBS, Invitrogen) and penicillin/streptomycin (Invitrogen) and maintained in a humidified incubator of 5% CO2 at 37C.

### Targeted image-based single-cell spatial transcriptomics (scST)

#### scST probe panel design

The scST probe panel was designed to include 75 CSP candidate genes described by Apostolo et al. (2020)^18^, complemented with markers for major brain cell types. Specifically, the panel contained markers for neurons (4), excitatory neurons (2), inhibitory neurons (2), astrocytes (5), microglia (2), oligodendrocytes (2), and endothelial cells (2). To enable potential discrimination of hippocampal astrocyte region-specific subtypes, an additional six markers associated with hippocampal astrocyte subpopulations were included based on Batiuk et al. 2020^20^; however, these markers did not allow reliable astrocyte subtype resolution in the final dataset. The resulting 100-plex target gene probe library (**Suppl st-panel.xlsx**) was designed and synthesized by Resolve Biosciences for use with the Molecular Cartography platform.

#### scST sample preparation

Three postnatal day 28 (P28) C57BL/6J male wild-type mice were anesthetized with isoflurane and euthanized by decapitation. Brains were rapidly extracted and immediately frozen in tissue embedding molds (Polysciences, #18985) containing FSC 22 compound (Leica Biosystems, #3801480) by immersion in isopentane cooled to -60 to -70 °C on dry ice. Samples were stored at -80 °C overnight prior to sectioning. Coronal hippocampal cryosections (10 µm) were cut using a Leica CM3050 cryostat. For each mouse, two sections were collected intermittently (every other section), with intervening sections discarded to ensure non-overlapping sampling along the rostro–caudal axis. Sections were trimmed to isolate the hippocampus and mounted within the capture areas of prechilled Resolve Biosciences slides. Mounted sections were stored at -80 °C for 1-3 days and then shipped overnight on dry ice to Resolve Biosciences in Monheim am Rhein, Germany where the Molecular Cartography protocol was performed. In total, two technical replicates (sections) per brain, from 3 brains were shipped. Regions of interest (ROIs) were selected for each section based on a brightfield overview scan.

#### scST raw data processing

Resolve Biosciences provided raw DAPI images and corresponding transcript coordinate tables for each imaged region of interest (ROI or sample). Of the six hippocampal samples, one was excluded from further analysis because it did not include the complete hippocampal structure.

Data processing was performed in Python using Jupyter Notebooks within a Conda environment on a high-performance computing (HPC) cluster, with core packages SpatialData (v0.6.1^54^) and Harpy (v0.2.0^55^). Transcript coordinate tables and raw DAPI images from each sample were first imported into a SpatialData object. To reduce background signal in the DAPI images, intensity values were min-max normalized, followed by contrast enhancement using contrast-limited adaptive histogram equalization (CLAHE) implemented via Harpy wrapper functions. DAPI-based cell segmentation was performed using the Cellpose-SAM model (v4.0.8^56^), executed through the Harpy’s wrapper function for cell segmentation and accelerated using an NVIDIA A100 GPU. Cellpose parameters were optimized iteratively and were the following: diameter 180 pixels, flow_threshold 0.9, cellprob_threshold -6 and min_size 160 pixels. Transcripts were allocated to their respective DAPI-segmented cell masks using the Harpy’s allocator function and the proportion of transcripts allocated to putative cells was assessed for each sample. The resulting SpatialData object was subsequently written to a Zarr store to ensure reproducibility and facilitate downstream analyses.

#### scST downstream data analysis

Downstream analyses were performed in Python using Jupyter Notebooks within a Conda environment on a high-performance computing (HPC) cluster. Core packages included SpatialData (v0.6.1^54^), AnnData (v0.12.7^57^), Harpy (v0.2.0^55^), Scanpy (v1.11.5^58^) and scVI (v1.4.1^59^). The processed SpatialData object was loaded from Zarr for further analysis.

Cells with fewer than 10 detected transcripts were excluded, and genes detected in fewer than 5 cells per sample were removed using Harpy’s preprocess_transcriptomics function. After filtering, 50.640 cells and 97 genes were retained. Given the limited size of the gene panel, highly variable gene selection was not performed. A merged AnnData object was constructed using Scanpy’s concatenation functionality, excluding per-sample outputs generated by preprocess_transcriptomics

Batch correction and latent embedding was performed using scVI. Prior to model training, the AnnData expression matrix was reset to raw counts. The scVI model was trained on the filtered dataset, incorporating segmented cell size as a covariate. Model training was accelerated using an NVIDIA A100 GPU and convergence was manually assessed. The learned latent representation was used to compute nearest neighbors with 40 nearest neighbors. Dimensionality reduction was performed using Uniform Manifold Approximation and Projection (UMAP) as implemented in Scanpy, using default parameters. Cells were clustered using the Leiden algorithm with a resolution parameter of 0.5, yielding 15 clusters with robust cross-sample integration.

Cluster assignments were subsequently projected back into the spatial context for each sample. Marker genes for each cluster were identified using Scanpy’s rank_genes_groups function using the Wilcoxon rank-sum test. Cell type annotation was performed manually based on spatial cluster localization patterns and cluster-specific marker gene expression. Annotated cell type labels were transferred back to the aggregated SpatialData object and distinct colors were assigned to each cell type. The trained scVI model was saved and the updated SpatialData object were saved to Zarr stores for reproducibility.

Quality control (QC) visualizations of scST analysis included: (i) dot plot of cell types with cell type markers, (ii) spatial plots of cell types per sample and across all samples, (iii) UMAP visualization colored by cell types and samples, (iv) distributions of genes per cell and counts per cell across cell types and samples; (v) numbers of cells per cell type and per sample; and (vi) distributions of cell types across samples.

Differential expression analysis was performed using scVI’s differential expression functionality with batch correction, filtering outlier cells, mode as change and weights as uniform. Comparisons of the following were made: Astrocytes vs CA1 Neurons, and vs CA2/CA3 Neurons, and vs DG Granule Neurons. For each comparison, differentially expressed genes (DEGs) were selected using an false discovery rate (FDR) threshold of 0.05 and an absolute log fold change greater than 1. Cell type marker genes were excluded from downstream interpretation. All astrocytic DEGs and the top three neuronal DEGs per comparison were retained. Log fold changes of candidate genes were visualized using heatmaps generated with Seaborn (v0.13.2^60^), and gene expression patterns were further visualized using Scanpy dot plots.

### CSP candidate profiling

#### Sample preparation for immunohistochemistry (IHC)

At P28, C57Bl/6J wildtype mice were anesthetized with a lethal dose of anesthetics (1 μl/g xylazine (VMB Xyl-M 2%), 2 μl/g ketamine (Eurovet Nimatek, 100 mg/ml), and 3 μl/g 0.9% saline) and checked for reflexes. Once all reflexes subsided, the abdominal and thoracic cavities were opened, and animals were transcardially perfused using a gravity-based system for 2 min with ice-cold PBS, followed by 7 min with ice-cold 4% PFA in PBS. Brains were gently removed from the skull and vertebrae, dissected out, and post-fixed ON in 4% PFA in PBS. Brains were then washed three times with PBS and cryoprotected in 30% sucrose in PBS at 4°C for at least two days (until brains sank). Subsequently, brains were frozen in a tissue embedding mold (Polysciences, #18985) in OCT compound (Sakura, #4583) by immersion in isopentane cooled to -60 to -70°C on dry ice. After at least one ON at -80°C, 20 μm coronal cryostat sections were cut using a Leica CM3050. Sections were collected intermittently (every other section), with intervening sections discarded, to ensure non-overlapping sampling along the rostro-caudal axis. Collected sections were mounted on slides and stored at -80°C until further processing.

#### IHC

Sections were equilibrated to RT for 20 min after removal from -80°C. Sections were then washed once in 1X PBS for 5 min and permeabilized using PBS-0.05% Triton X-100 for 20 min. Subsequently, sections were blocked for 2 h in blocking solution containing 10% NHS, 0.5M Glycine, 0.5% Triton X-100 in 1X PBS. When mouse primary antibodies were used, a donkey anti-mouse Fab fragment (Jackson ImmunoResearch, 715-007-003) was included in the blocking solution at a 1:50 dilution to block endogenous mouse immunoglobulins. Next, sections were washed either once or five times with PBS-0.05% Triton X-100, depending on whether a donkey anti-mouse Fab fragment was used. Primary antibodies (**Table S2**) were diluted in antibody solution (5% NHS, 0.5% Triton X-100, 0.02% gelatin in PBS) and incubated ON at 4°C. Next day, sections were washed for 10 min in PBS-0.05% Triton X-100 at least five times. Secondary antibodies (**Table S3**) were diluted in antibody solution and incubated for 2 h at RT. Sections were subsequently washed for at least 5 times with PBS-0.05% Triton X-100 and counterstained with DAPI for 10 min. After, sections were washed for two more times and mounted using Mowiol. Super-resolution images of synapses were acquired on a Zeiss LSM880 confocal microscope equipped with an Airyscan detector and a 63X oil-immersion objective (NA 1.4). All images were acquired in Airyscan mode under identical acquisition settings across experimental conditions to enable qualitative comparison.

#### smFISH (RNAscope) with IHC

Same preparation protocol was used as mentioned in IHC section with slight modifications. After ON post fixation, brains were embedded in agarose for 2 hr and sectioned at 40 μm using a vibratome (Campden Instruments 7000 smz 2). Free-floating sections were collected on slides and air dried in an oven at 40°C for 5 min. Then, smFISH was performed according to manufacturer instructions (RNAscope multiplex v2 kit, ACD #323110). Briefly, sections were treated for 10 min with a hydrogen peroxide solution and 20 min with protease IV, with 1X PBS washing in between. RNAscope probes were hybridized at 40°C for 2 hours, and subsequently, a horseradish peroxidase signal was developed. Gpr37l1, Hepacam, Vcam1, Alcam, Cadm4 and DapB negative control probes were used in the C1-channel with Opal dye 570 (**Table S4**). After smFISH, sections were blocked for 2 h in blocking solution containing 10% NHS, 0.5M Glycine, 0.5% Triton X-100 in 1X PBS. Next, sections were immuno-stained with S100b as an astrocyte marker, VGLUT1 to delineate the hippocampal layers, DAPI as nuclear staining and mounted using Mowoil as in the IHC section. Confocal Images were acquired using a Zeiss LSM880 confocal laser scanning microscope using the 10X or 20X objective. All imaging was performed in confocal mode under identical acquisition settings, to enable qualitative comparison.

#### Synaptogliosome preparation

Synaptogliosomes were prepared according to Mazare et al. 2020^22^ with slight modifications. At P28, C57Bl/6J wildtype mice were anesthetized with isoflurane and euthanized by decapitation. Brains were removed, and hippocampi were rapidly dissected in ice-cold Hank’s Balanced Salt Solution (HBSS, Gibco, 14175-095). All steps were performed on ice. Tissue was transferred to a pre-cooled 1 mL Dounce homogenizer, residual HBSS was aspirated, and replaced with 800 μl homogenization buffer (0.32 M sucrose, 10 mM HEPES, 2 mM EDTA supplemented with protease inhibitor cocktail (Roche, 11697498001), pH adjusted to 7.4) per brain (2 hippocampi). 1 brain was considered as one biological replicate. Homogenization was performed by 20 gentle strokes with pestle A (narrow) followed by 20 strokes with pestle B (wide), ensuring minimal foaming and avoiding air bubbles.

The hippocampal homogenate was transferred to pre-cooled 1 mL eppendorfs and centrifuged at 900 x g for 15 min at 4°C to pellet the nuclear and debris fraction (P1). The resulting supernatant (S1) was collected and centrifuged at 16,000 x g for 15 min 4°C. The supernatant (S2) was discarded, and the pellet (P2) was resuspended in 600 μl of homogenization buffer and centrifuged again at 16,000 x g for 15 min 4°C. The final P3 pellet contained the synaptogliosomes and was resuspended in 300 μl of 1:1 mixture of homogenization buffer and 2X lysis/wash buffer (50 mM Tris HCl, 300 mM NaCl, 2 mM EDTA, 2% NP-40, 10% glycerol, supplemented with protease inhibitor cocktail (Roche, 11697498001), pH 7.4).

Protein was extracted by rotating the samples at 4°C for 1 h, and the protein concentration was determined using the Pierce BCA protein assay kit (Thermo Fisher, 23227) according to the manufacturer’s instructions.

#### Sample preparation for developmental timepoints

Littermate pups were used for hippocampal lysates at P1, P7, P14, P21 and P28. Briefly, pups were anesthetized with isoflurane and euthanized by decapitation. Brains were removed, and hippocampi were rapidly dissected in ice-cold Hank’s Balanced Salt Solution (HBSS, Gibco, 14175-095). All steps were performed on ice. Tissue was transferred to a pre-cooled 1 mL Dounce homogenizer, residual HBSS was aspirated, and replaced with extraction and lysis buffer (25 mM Tris HCL pH 7.5, 150 mM NaCl, 1 mM EDTA, 1% NP-40, 0.5% sodium deoxycholate, 0.1% SDS supplemented with protease inhibitor cocktail (Roche, 11697498001), pH adjusted to 7.4) per brain (2 hippocampi). Volume of extraction and lysis buffer for homogenization was adjusted according to age: P1: 0.2 mL, P7: 0.2 mL, P14: 0.5 mL, P21: 1 mL, P28: 1 mL. Homogenization was performed by 20 gentle strokes with pestle A followed by 20 strokes with pestle B, ensuring minimal foaming and avoiding air bubbles. Protein was extracted by rotating the samples at 4°C for 30 min, followed by centrifugation at 14,000 rpm at 4°C for 10 min. Supernatant was collected and protein concentration was determined using the Pierce BCA protein assay kit (Thermo Scientific, 23227) according to the manufacturer’s instructions. Aliquots were made and stored at -80 °C until further processing.

#### Western blot

Hippocampal lysates from P1, P7, P14, P21 and P28 (20 μg) were mixed with 5X SDS loading buffer brought to volume with distilled water, and boiled for 10 min at 95 °C. Proteins were separated on a 4-20% precast polyacrylamide gel (Bio-Rad, 4561094) at 100V for ∼1.5 h and transferred to a 0.2 µm nitrocellulose membrane using a Trans-Blot Turbo Transfer System (Bio-Rad, 1704150EDU) under the mixed-molecular weight program. Membranes were stained with the Revert™ Total Protein Stain Kit (Li-COR, 926-11021) according to the manufacturer’s protocol.

Membranes were blocked in 5% milk in TBST buffer (25 mM tris-base pH 7.5, 300 mM NaCl, and 0.05% Tween 20) for 1 h at RT. Primary antibodies (**Table S2**) were incubated overnight at 4°C on a rocking platform, and secondary antibodies (**Table S3**) were incubated for 1 h at RT on a rocking platform. After each antibody incubation, membranes were washed four times with TBST buffer. Chemiluminescent signals were detected using the SuperSignal West Femto Maximum Sensitivity Substrate (Thermo Fisher, 34094) and imaged on an Amersham ImageQuant 800 (Cytiva). Images were quantified by measuring band intensities in ImageJ using the Gels-Plot lanes tool. Protein expression levels were first normalized to total protein levels and then to P28 levels. Data was plotted using Seaborn’s (v0.3.12) lineplot function.

### CRISPR/Cas9 *in-vitro* KO validation

#### Low-titer AAV production

Low-titer adeno-associated virus (AAV) was produced from cell culture supernatant. HEK cells were seeded in a 12-well plate and transfected at ∼80% confluency using polyethylenimine (PEI). Each well received a DNA:PEI complex containing 800 ng pDelta F6 plasmid, 400 ng RepCap 2/5 plasmid, and 400 ng vector transgene plasmid of interest (AAV-Lck-GFP, AAV-LacZ-gRNA#1 and #2, AAV-VCAM-gRNA#1, #2 and #3), combined with 5.7 μl PEI in a total volume of 100 μl OptiMEM (Thermo Fisher Scientific, #31985062). Culture supernatants were harvested 3 days after transfection, clarified by filtration through a 0.45-μm syringe filter, and concentrated by sequential centrifugation using a 100 kDa molecular weight cutoff Amicon Ultra-15 centrifugal filter unit (Millipore, UFC9100) to a final volume of ∼40 μl.

#### Primary hippocampal astrocyte cultures and transduction

Primary hippocampal astrocyte cultures were prepared from dissected P1 C57Bl/6J or H11-Cas9 mice from both sexes. Briefly, hippocampi were dissected in HBSS (Gibco, 14175-095) and enzymatically digested in 0.25% trypsin-EDTA (Gibco, 15090-046) in HBSS with 10 mg/mL DNase I (Sigma, Dn25-100mg) at 37°C for 15 min with occasional swirling. Then, tissue was washed three times with glial culture medium (500 mL): 430 mL MEM (Gibco, 31095-029), 10% FBS (Gibco, A5256701), 15 mL 20% Glucose (homemade), 5 mL Pen/Strep (Gibco, 15140-122). Next, the tissue was mechanically dissociated by gentle trituration with a 10 mL glass pipet for a maximum of 15 times, incubated for 10 min at 37°C and again dissociated with a 5 mL glass pipet for a maximum of 15 times. Then, the cell suspension was passed through a 70 μm cell restrainer and washed two times with fresh glial medium.

After centrifugation for 5 min at 1000 rpm, the cell pellet was resuspended in glial medium and cell density was determined. Approximately 7.5 x 10^6^ cells were plated per 75 mm^2^ flask and maintained in glial medium. To enrich for astrocytes, non-astrocyte cells were removed by forcefully manual shaking the flasks for 15s on DIV1 and DIV4, leaving a confluent astrocyte monolayer. At DIV7, astrocytes were harvested and replated into 6-well plates at ∼30 x 10^5^ cells per well. Half of glial medium was replaced every 3-4 days. At DIV10, 10 μM of 5-fluoro-2’-deoxyuridine was added to limit proliferative overgrowth.

For the VCAM1 and GPR37L1 protein level experiments (**Fig. S4A**), astrocytes at DIV14 were washed twice with ice cold TBS containing 1 mM CaCl2 and 1 mM MgCl2, followed by incubation on ice for 20 minutes in 300-700 μl/well membrane solubilization buffer (25 mM Tris pH7.4, 150 mM NaCl, 1 mM Cacl2, 1 mM MgCl2, 0.5% NP-40, and protease inhibitors) with gentle agitation. Astrocyte lysates were collected and pooled, briefly vortexed, and centrifuged at 15.000 x g for 10 min at 4°C to pellet non-solubilized material. The supernatants were collected, and protein concentration was determined using the Pierce^TM^ BCA protein assay kit (Thermo Scientific, #23227) according to the manufacturer’s instructions. Aliquots were prepared and stored at 80°C. For the VCAM1-gRNA efficiency experiments (**Fig. S4B**), hippocampal astrocytes were transduced at DIV14 with 35 μl of low-titer AAVs (∼1:60 dilution). At DIV28, the astrocyte protein lysates were collected using the same procedure as described above. Notably, in these in vitro experiments, only 60 to 80% of the astrocytes were estimated to be infected with AAV-GFP virus by eye, suggesting a similar infection rate for AAV-gRNA viruses. The partial transduction of astrocytes likely explains the residual VCAM1 protein levels in Vcam1 gRNA-infected cultures.

#### Western blot

Astrocyte lysates (5-10 μg) were mixed with 5X SDS loading buffer brought to volume with distilled water, and boiled for 45 min at 45 °C. Proteins were separated on a 4-20% precast polyacrylamide gel (Bio-Rad, 4561094) at 100V for ∼1.5 h and transferred to a 0.2 µm nitrocellulose membrane using a Trans-Blot® Turbo™ Transfer System (Bio-Rad, 1704150EDU) under the mixed-molecular weight program. Membranes were stained with the Revert™ Total Protein Stain Kit (Li-COR, 926-11021) according to the manufacturer’s protocol.

Membranes were blocked in 5% milk in TBST buffer (25 mM tris-base pH 7.5, 300 mM NaCl, and 0.05% Tween 20) for 1 h at RT. Primary antibodies (**Table S2**) were incubated overnight at 4°C on a rocking platform, and secondary antibodies (**Table S3**) were incubated for 1 h at RT on a rocking platform. After each antibody incubation, membranes were washed four times with TBST buffer. Chemiluminescent signals were detected using the SuperSignal™ West Femto Maximum Sensitivity Substrate (Thermo Fisher Scientific, 34094) and imaged on an Amersham™ ImageQuant™ 800 (Cytiva).

#### Western blot data analysis

For the VCAM1-gRNA efficiency experiments, band intensities of the total protein staining and VCAM1 protein levels were analyzed using Fiji (ImageJ). VCAM1 signal intensities were first normalized to total protein levels for each lane, and normalized values were subsequently expressed relative to the LacZ-gRNA#1 control within each experiment. Statistical significance was evaluated using analysis of variance (ANOVA) with gRNA and experiment included as factors, followed by Dunn’s post hoc comparisons of each gRNA condition against the LacZ-gRNA#1 control, with Benjamini-Hochberg correction applied to control the FDR.

### CRISPR/Cas9 KO *in-vivo* synapse analysis

#### High-titer AAV production

High-titer AAV was produced and purified as previously described^61^. Briefly, six plates of 80% confluent HEK cells were transfected using PEI with 20 μg of pDelta F6 plasmid, 10 μg of RepCap 2/5 plasmid and 10 μg vector transgene plasmid of interest (AAV-LacZ-gRNA#1, AAV-VCAM-gRNA#2 and AAV-GPR37L1-gRNA) per plate. Following transfection, cells were maintained in DMEM supplemented with 5% FBS for 3 days.

Next, cells were harvested by scraping, spun at 1000 g for 10 mins at 4°C and pellets were lysed in lysis buffer (150 mM NaCl and 50 mM Tris HCl (pH 8.5)). Lysates were subjected to three freeze-thaw cycles, followed by centrifugation at 4000 x g for 15 mins at 4°C. Benzonase nuclease (Sigma-Aldrich, E1014) added at a concentration of 50 U/mL to supernatants for 30 mins at 37°C to remove contaminating cellular DNA. Lysates were subsequently centrifuged at 4000 x g for 10 min at RT.

Lysates were then filtered through a 0.45 μm filter and loaded onto an OptiPrep iodixanol gradient (15%, 25%, 40%, and 60%, Millipore, D1556) and ultracentrifuged at 300.000 x g for 100 min at 12°C. AAVs particles were carefully collected with an 18-gauge needle (Beckman Coulter) from between the 40% and 60% layers, and desalted and concentrated by centrifugation at 4000 x g for 30 min at 20°C in a prerinsed 100 kDa molecular weight cutoff Amicon Ultra-4 centrifugal filter unit (Millipore, UFC8100). AAVs were aliquoted and stored at -80°C.

AAV purity was assessed by silver staining using the ProteoSilver Silver Stain Kit (Sigma PROTSIL1-1KT). The AAV titers were determined by qPCR using LightCycler 480 SYBR Green I Master (Roche 04707516001) to normalize titers across viral preparations.

#### AAV stereotactic injections

Prior to surgery, P7 H11-Cas9 mouse mice were anesthetized with 5% isoflurane and positioned in a stereotact frame (Kopf Instruments) on a 37°C heated platform. Anesthesia was maintained with 2% isoflurane throughout the procedure. After disinfecting the mouse’s scalp, local anesthesia was administered by a subcutaneous injection with 100 μl of lidocaine (1% xylocaine). A midline incision was made to expose the skull.

To target the hippocampus at P7, stereotactic coordinates relative from lambda were: mediolateral (ML), 2.3 mm; anteroposterior (AP), 0.9 mm; dorsoventral (DV), 2.3 mm. 200 nL of AAV-U6::VCAM1- or AAV-U6::GPR37L1-gRNA-CAG::FLEX-Lck-GFP virus was injected ipsilateral hemisphere and AAV-U6::LacZ-gRNA-CAG::FLEX-Lck-GFP was injected in the contralateral hemisphere using a Nanoject III (Drummond Scientific) equipped with a beveled glass capillary at 4 nl/s. After 2 min, the capillary was slowly pulled out at ∼0.1 mm per 5 s to minimize backflow. The incision was closed using surgical glue (Millpledge Veterinary). Animals were monitored during recovery, and buprenorphine (0.1 mg/kg) was administered 6 h post-surgery for analgesia. All gRNA AAVs injected were at a concentration of ∼10^13^ GC/mL.

#### Immunohistochemistry sample preparation

At P21, mice were anesthetized with a lethal dose of anesthetics (1 μl/g xylazine (VMB Xyl-M 2%), 2 μl/g ketamine (Eurovet Nimatek, 100 mg/ml), and 3 μl/g 0.9% saline) and checked for reflexes. Once all reflexes subsided, the abdominal and thoracic cavities were opened, and animals were transcardially perfused using a gravity-based system for 2 min with ice-cold PBS, followed by 7 min with ice-cold 4% PFA in PBS. Brains were gently removed from the skull and vertebrae, dissected out, and post-fixed ON in 4% PFA in PBS. Brains were then washed three times with PBS and cryoprotected in 30% sucrose in PBS at 4°C for at least two days (until brains sank). Subsequently, brains were frozen in a tissue embedding mold (Polysciences, #18985) in OCT compound (Sakura, #4583) by immersion in isopentane cooled to -60 to -70°C on dry ice. After at least one ON at -80°C, 20 μm coronal cryostat sections were cut using a Leica CM3050. Sections were collected intermittently (every other section), with intervening sections discarded, to ensure non-overlapping sampling along the rostro-caudal axis. Collected sections were mounted on slides and stored at -80°C until further processing.

#### Immunohistochemistry (IHC)

Sections were equilibrated to RT for 20 min after removal from −80°C. Sections were then washed once in 1X PBS for 5 min and permeabilized using PBS-0.05% Triton X-100 for 20 min. Subsequently, sections were blocked for 2 h in blocking solution containing 10% NHS, 0.5M Glycine, 0.5% Triton X-100 in 1X PBS. When mouse primary antibodies were used, a donkey anti-mouse Fab fragment (Jackson ImmunoResearch, 715-007-003) was included in the blocking solution at a 1:50 dilution to block endogenous mouse immunoglobulins. Next, sections were washed either once or five times with PBS-0.05% Triton X-100, depending on whether a donkey anti-mouse Fab fragment was used. Primary antibodies (**Table S2**) were diluted in antibody solution (5% NHS, 0.5% Triton X-100, 0.02% gelatin in PBS) and incubated ON at 4°C. Next day, sections were washed for 10 min in PBS-0.05% Triton X-100 at least five times. Secondary antibodies (**Table S3**) were diluted in antibody solution and incubated for 2 h at RT. Sections were subsequently washed for at least 5 times with PBS-0.05% Triton X-100 and counterstained with DAPI for 10 min. After, sections were washed for two more times and mounted using Mowiol.

#### Super-resolution image acquisition

Super-resolution images of synapses were acquired on a Zeiss LSM880 confocal microscope equipped with an Airyscan detector and a 63X oil-immersion objective (NA 1.4). All images were acquired in Airyscan mode under identical acquisition settings across experimental conditions to enable quantitative comparison. For each field of view (FOV), a short Z-stack was acquired, and a single optical plane was selected using a predefined and standardized criterion applied uniformly across all samples. For each FOV, a single optical plane of 1032 x 1032 pixels was acquired at a zoom factor of 3X, corresponding to a pixel size of 40 nm x 40 nm, for both presynaptic (488 nm) and postsynaptic (647 nm) channels at 16-bit depth. The Airyscan detector was aligned prior to imaging each new slide. One FOV was acquired per section for each hippocampal lamina and each hippocampal hemisphere, with three sections sampled per brain, with four brains in total. To minimize selection bias, only DAPI and VCAM1 or GPR37L1 staining were used for the initial localization. This ensured that VCAM1 or GPR37L1 signal was depleted in the gRNA-injected hemisphere and preserved in the LacZ-gRNA-injected hemisphere prior to image acquisition. Presynaptic and postsynaptic channels were not consulted during field selection. Image acquisition was not performed blinded to experimental condition. Airyscan processing was performed directly on the microscope workstation immediately after the acquisition using a standard super-resolution processing parameter strength of 5. All images were stored in native ZEISS *CZI* format.

#### Custom automated image processing pipeline

A custom automated synapse processing and analysis pipeline was developed in Python (v3.11) and orchestrated using Nextflow (v25.04.8^62^) to analyze super-resolution Airyscan synapse images. The pipeline uses a small custom-made Python library for synapse analysis, developed based on scikit-image (v0.25.2^63^) functions and published on PyPI (synapse-counting, v0.1.3). Images were imported using the czifile library (v2019.7.2.1). Preprocessing parameters and parameter ranges for local peak maxima detection were empirically optimized per synapse type (VGLUT1-PSD95 and VGAT-GEPH) and per hippocampal lamina using custom Jupyter notebooks prior to pipeline execution and stored as *JSON* configuration files, applied uniformly across all analyses. Image files followed a standardized naming convention encoding experimental metadata, including experimental batch, brain and section identifiers, gRNA condition, hippocampal region, and hippocampal lamina. Filename segments were automatically parsed to extract experimental condition, brain identifier, and hippocampal lamina, enabling automated grouping and downstream analysis.

Depending on lamina-specific signal characteristics, preprocessing steps could include background subtraction using a rolling-ball algorithm, Gaussian smoothing, contrast enhancement using contrast-limited adaptive histogram equalization, top-hat filtering, image binarization using a predefined auto thresholding algorithm, watershed-based segmentation, and exclusion of small objects below a minimum puncta size threshold to remove noise. Again, these parameters ares stored in a *JSON* configuration file in the GitHub code repo (please see section code availabity).

Synaptic colocalization was quantified using four complementary metrics: Manders’ overlap coefficient (measure.manders_overlap_coeff), Pearson’s correlation coefficient (measure.pearson_corr_coeff function), binary pixel overlap following image binarization (counting pixels using numpy), and spatial colocalization of local intensity maxima (feature.peak_local_max). For local maxima-based colocalization, presynaptic and postsynaptic puncta were identified as local intensity peaks, and synapses were defined when corresponding peaks were located within a specified maximum Euclidean distance.

To minimize bias in puncta detection, parameters governing local peak identification and synapse assignment were optimized separately for each batch of presynaptic and postsynaptic images per hippocampal lamina. Optimized parameters included intensity thresholds for detecting presynaptic and postsynaptic local peaks, the maximum allowable distance between presynaptic and postsynaptic peaks, and the minimum inter-peak distances for presynaptic and postsynaptic puncta. Parameter optimization was performed using the Optuna library (v4.3.0^64^), based on a Tree-structured Parzen Estimator. Optimization was performed exclusively on LacZ-gRNA control images by maximizing an objective function defined as the mean scaled difference in colocalization between original images and spatially 90° rotated control images, normalized by the number of images. This approach provided an internal negative control for random colocalization. The local peak optimization procedure was repeated 100 times per synapse type and per hippocampal lamina.

In addition to colocalization metrics, puncta-based synaptic properties were quantified separately for pre- and postsynaptic markers, including mean fluorescence intensity, puncta density normalized per 100 µm², total staining area, and mean puncta size.

All Python-based image processing steps were parallelized at the image level using Dask (v2025.2.0^65^), while workflow-level parallelization and data aggregation were handled by Nextflow. Each analysis process produced structured output files in *CSV* format, which were subsequently aggregated, cleaned, analyzed, and visualized using standard Python libraries. The pipeline was executed on high-performance computing infrastructure using SLURM and run independently for each combination of synaptic markers. All pipeline parameters, including preprocessing settings, optimization ranges, image channel assignments, filename parsing rules, and execution environment, were specified in centralized configuration files to ensure reproducibility. The pipeline can be executed using Conda^66^, Docker^67^, or Apptainer^68^ containers.

#### Downstream data analysis

The metrics results outputs from the custom pipeline were loaded into R (v4.5.0) running in a custom Docker development container (v28.1.1^67^) to ensure computational reproducibility and platform independence. Python-based (v3.11) analyses were performed in Conda-managed environments (v25.1.1^66^).

First, to account for brain-to-brain variability prior to multivariate analysis, synaptic metrics were analyzed using linear-mixed effects modeling. Specifically, each synaptic metric was iteratively modeled, after tidyverse (v2.0.0) data wrangling, across hippocampal laminae using a linear mixed-effects model with a random intercept for brain (metric ∼ 1 + (1 ∣ Brain)) implemented in lme4 (v1.1-37^69^). Residuals from these models were assembled into a residual matrix across metrics (one row per sample) and exported as a *CSV* file. PCA was then performed on the standardized residual matrix using prcomp (stats v3.6.2) to assess hemisphere variation in synaptic metrics by gRNA condition after accounting for brain-level effects. The two first principal components were visualized using ggplot2 (v3.4.3). The residual matrix was additionally imported into Python, where values were scaled using StandardScaler from scikit-learn (v1.6.1). To identify and visualize clustering of synaptic metrics across brains and hemispheres, hierarchical clustering was performed on the scaled residuals using clustermap from the seaborn (v0.13.2) with correlation used as distance metric.

Second, to formally test for differences in synaptic metrics between LacZ-gRNA- and VCAM1-gRNA- or GPR37L1-gRNA-injected hemispheres, synaptic metrics were analyzed using a linear-mixed effects model to fit the raw metric values for each synapse type (VGLUT1–PSD95 or VGAT–GEPH. For each metric, data were extracted along with gRNA condition, hippocampal lamina, section identifier, and brain identifier using tidyverse (v2.0.0). Metrics were log2-transformed after addition of a pseudocount (log2 (x + 1)) to improve normality. For each metric, a linear mixed-effects model was fit with gRNA condition and hippocampal lamina as fixed effects, including their interaction, and brain included as a random intercept to account for repeated measurements within brains: metric ∼ gRNA × hippocampal lamina + (1 ∣ Brain). Models were fit using restricted maximum likelihood (REML). Model assumptions were evaluated using visual inspection of residual Q-Q plots and residuals versus fitted values. Post hoc comparisons between gRNA conditions were performed within each hippocampal lamina using estimated marginal means using the emmeans package (v1.11.1). Pairwise contrasts were adjusted for multiple testing using false discovery rate (FDR) correction across each metric. For each comparison, estimated marginal means for each gRNA condition and the corresponding contrast statistics were reported. All analyses were performed independently for each synaptic metric across hippocampal laminae, and results were aggregated across metrics for downstream interpretation. The resulting matric was imported into Python. Log2 fold changes comparing LacZ-gRNA- with VCAM1-gRNA- or GPR37L1-gRNA-injected hemisphere were visualized as heatmaps using the heatmap function from seaborn (v0.13.2). Numpy (v2.2.5) and pandas (v2.2.3) were used for data wrangling.

### Synaptosome co-immunoprecipitation coupled with mass spectrometry (MS)

#### Crude synaptosome preparation

P21 C57Bl/6J wildtype mice were anesthetized with isoflurane and euthanized by decapitation. Brains were removed, and hippocampi were rapidly dissected in ice-cold Hank’s Balanced Salt Solution (HBSS, Gibco, 14175-095). All steps were performed on ice. Tissue was transferred to a pre-cooled 2 mL Dounce homogenizer, residual HBSS was aspirated, and replaced with 2 mL homogenization buffer (0.32 M sucrose, 10 mM HEPES, 2 mM EDTA supplemented with protease inhibitor cocktail (Roche, 11697498001), pH adjusted to 7.4). Four hippocampi from two brains (one male and one female) were pooled and processed as one biological replicate. Homogenization was performed by 20 gentle strokes with pestle A followed by 20 strokes with pestle B, ensuring minimal foaming and avoiding air bubbles.

The hippocampal homogenate was transferred to pre-cooled 2 mL tubes (Fisherbrand, 11558252) and centrifuged at 1,000 x g for 10 min at 4°C to pellet the nuclear and debris fraction (P1). The resulting supernatant (S1) was collected and centrifuged at 14,000 x g for 15 min 4°C. The supernatant (S2) was discarded, and the pellet (P2), containing crude synaptosomes, was resuspended in 1 or 2 mL of a 1:1 mixture of homogenization buffer and 2X lysis/wash buffer (50 mM Tris HCl, 300 mM NaCl, 2 mM EDTA, 2% NP-40, 10% glycerol, supplemented with protease inhibitor cocktail (Roche, 11697498001), pH 7.4).

Protein was extracted by rotating the samples at 4°C for 1 h, and the protein concentration was determined using the Pierce BCA protein assay kit (Thermo Scientific, 23227) according to the manufacturer’s instructions.

#### Co-immunoprecipitation

For each biological replicate, the crude synaptosomal sample was split into two equal fractions to enable paired processing for immunoprecipitation (IP) and IgG control conditions. Each fraction (500 or 600 µg total protein) was prepared in 1 mL of 1X lysis/wash buffer in 1.5 mL low-binding tubes (Eppendorf, 0030108442).

To pre-clear the samples, 25 µl of pre-washed Dynabeads protein G (Invitrogen, 10004D), washed three times with 1× lysis/wash buffer, were added and incubated at 4°C for 1 h with rotation. After incubation, the unbound fraction (supernatant) was transferred to a new low-binding tube. Next, 10 µg of antibody (**Table S1**) and 50 µl of pre-washed Dynabeads protein G were added to the pre-cleared samples and incubated overnight at 4°C with rotation.

The following day, beads were collected using a DynaMag magnetic rack (Invitrogen, 12321D), and the unbound fraction was removed. Beads were washed three times with 1 mL of 1X lysis/wash buffer, mixing gently for 2 min per wash. After each wash, the beads were collected on the magnetic rack for 2 min before the buffer was removed.

#### MS sample preparation

Proteins were eluted from the beads and digested into peptides using the S-Trap Micro Spin Columns (Protifi, C02-MICRO-80) according to the manufacturer’s instructions. Briefly, beads were resuspended in 50 µl of 5% SDS, 50 mM triethylammonium bicarbonate (TEAB, pH 8.5), boiled for 5 min at 95 °C, and cooled to RT. Using a magnetic rack, beads were separated, and supernatants were transferred to a new low-binding tubes. Eluates were further reduced with 5 mM Tris (2-carboxylethyl) phosphine (TCEP) buffer at 55 °C for 15 min and subsequently alkylated with 20 mM iodoacetamide (IAA) at RT for 15 min in the dark. Samples were acidified with 5.4 µl of 27.5% phosphoric acid (v/v).

Samples were mixed with 360 µl of 90% methanol in 100 mM TEAB and loaded onto the S-Trap Micro Spin Column trap the proteins and further centrifuged at 4000 x g for 1 min. Columns were then washed three times with 300 µl of 90% methanol in 100 mM TEAB.

Proteins were digested on-column with 1 µg of trypsin in 20 µL of 50 mM TEAB at 37 °C overnight to ensure complete protein digestion. Peptides were sequentially eluted from the column at 4000 x g for 1 min using 40 µL of each of the following buffers: (i) 50 mM TEAB (pH 8.5); (ii) 0.2% formic acid in LC-MS grade H_2_O; and (iii) 50% acetonitrile in LC-MS grade H_2_O. Eluates were pooled and completely dried using a CentriVap® SpeedVac™ vacuum concentrator (Labconco).

#### Western blot quality control of immunoprecipitation fractions

A 15 µL aliquot (1.5 % input) was collected before sample splitting (into IP and IgG) and after addition of 5% SDS (for IgG control and IP). Samples were mixed with 5X SDS loading buffer and boiled for 5min at 95 °C (input only). Proteins were separated on a 4-20% precast polyacrylamide gel (Bio-Rad, 4561094) at 100V for ∼1.5 h and transferred to a 0.2 µm nitrocellulose membrane using a Trans-Blot® Turbo™ Transfer System (Bio-Rad, 1704150EDU) under the mixed-molecular weight program.

Membranes were blocked in 5% milk in TBST buffer (25 mM tris-base pH 7.5, 300 mM NaCl, and 0.05% Tween 20) for 1 h at RT. Primary antibodies (**Table S2**) were incubated overnight at 4 °C on a rocking platform, and secondary antibodies were incubated for 1 h at RT on a rocking platform. After each antibody incubation, membranes were washed four times with TBST buffer. Chemiluminescent signals were detected using the SuperSignal West Pico PLUS Chemiluminescent Substrate (Thermo Fisher Scientific, 34580) and imaged on an Amersham ImageQuant 800 (Cytiva).

GPR37L1 protein migrates at a molecular weight overlapping with IgG fragments, thus, conventional detection using standard secondary antibodies was unsuccessful. The Mouse TrueBlot ULTRA kit (Rockland, 88-8887-31) was subsequently tested to minimize IgG interference, however, detection remained unsuccessful, likely due to low sensitivity of the secondary antibody toward the primary mouse IgG_2_A.

#### Mass spectrometry

Tryptic peptides were reconstituted in 20 µL of 0.5% acetonitrile (ACN) and 99.5% water containing 0.1% formic acid (FA). From this, 2 µL of the sample was loaded onto an Evotip following the manufacturer’s guidelines for analysis using an Evosep One system coupled with a ZenoTOF 7600 mass spectrometer (AbSCIEX), utilizing Sciex OS (version 3.3 or higher). The peptides were separated via the 30 samples-per-day (30SPD) method with a 44-minute gradient using EV1137 Performance column (15 cm x 150 µm, 1.5 µm) connected to the low microelectrode for flow rates of 1-10 μL/min. Mobile phases included 0.1% FA in LC–MS-grade water (buffer A) and ACN/0.1% FA (buffer B). The ZenoTOF mass spectrometer operated in SWATH mode with an OptiFlow Turbo V ion source using the following source settings: spray voltage of 4500 V, gas 1 at 20 psi, gas 2 at 10 psi, curtain gas at 40 psi, CAD gas at 7, temperature range of 100 °C, and column temperature of 40 °C. The Zeno SWATH DIA method consisted of 85 variable-width windows enabling the selection of precursors from 350-1250 m/z. TOF-MS scans ranged from 350–1250 m/z with an accumulation time of 25 ms, and TOF-MS/MS scans covered 230–1400 m/z with accumulation times of 20 ms with dynamic collision energy turned on, a charge state of 2, and Zeno pulsing enabled.

#### MS raw data processing

Raw data were exported as *WIFF* files and converted to *mzML* format using the msConvert tool from ProteoWizard^70^ within an Apptainer^68^ container. Converted files were analyzed using Data-Independent Acquisition by Neural Networks (DIA-NN) v2.2.0^71^ in library-free mode using an Apptainer container. First, a spectral library was generated from the *Mus musculus* reference proteome (Uniprot UP000000589, accessed 20250930). Next, *mzML* files were processed against the predicted library with identical parameters supplemented with common contaminants sequences (cRAP, accessed 20250304).

Trypsin specificity was defined as cleavage after lysine (K) and arginine (R), excluding cleavage before proline, allowing up to one missed cleavage. Methionine oxidation and N-terminal acetylation were set as variable modifications, with up to one variable modification per peptide. Peptide lengths were restricted to 7-30 amino acids. DIA-NN was set to perform N-terminal methionine excision, and report results at the peptidoform level. The precursor m/z range was restricted to 350-1250, and precursor charge states between +1 and +4 were considered. The MS2 and MS1 mass accuracies were set to 20 ppm and 12 ppm, respectively. Quantification was performed using extracted ion chromatograms (XICs) with retention time profiling. The protein-level false discovery rate (FDR) was controlled at 1% (q < 0.01) and protein inference was performed in relaxed mode. The neural network classifier was operated in double-pass mode.

All data processing was executed on a high-performance computing (HPC) cluster with parallel execution where possible, using containerized workflows to ensure full reproducibility and platform independence.

#### MS downstream data analysis

The protein group outputs from DIA-NN were loaded into R (v4.5.0) running in a custom Docker development container (v28.1.1^67^) to ensure reproducibility and platform independence. First, Immunoglobulin (IgG) proteins stemming from antibodies and human proteins were removed prior to analysis. Using dplyr (v1.1.3), only proteins detected in ≥4 replicates in at least one condition were retained. Data were log2 transformed and missing values were imputed using a left-censored deterministic minimal value approach with the MinDet function from the imputeLCM package (v2.1).

Principle components analysis (PCA) was performed using prcomp (stats v3.6.2). Differential protein enrichment between IP and control conditions was tested using Limma (v3.62.1), employing a paired design (∼replicate + condition) and empirical Bayes moderation. Significantly enriched proteins were defined as those with an Benjamini-Hochberg (BH) adjusted P-value < 0.05 and log2 fold change > 1. For rank plot visualization, log2-transformed ratios between IP (mean) and IgG (mean) intensities were computed for each protein, and proteins were ranked based on these values.

For cross-referencing with synaptic proteome datasets, data from Van Oostrum et al.^72^ were imported into R, while dataset from Sorokina, Mclean and Croning et al.^73^ were accessed using RSQLite (v2.3.1). UniProt subcellular localization annotations were obtained using the UniProtR package (v2.4.0). “Cell surface proteins (CSPs)” were retained if annotated as “cell membrane,” “secreted,” “extracellular space,” “extracellular matrix,” or “cell surface.” Proteins annotated as “peripheral membrane protein” or “mitochondrion membrane” were excluded to avoid intracellular membrane-associated or mitochondrial contaminants.

Gene Ontology (GO) enrichment of GPR37L1 co-IP proteins was performed with the enrichGO function from the clusterProfiler package (v4.14.6), using the Biological Process (BP) ontology. The mouse reference proteome was used as background. Significance thresholds were set at P-value < 0.05 and q < 0.05, adjusted using the BH correction method. GO term visualization was performed using the built-in plotting tools. SynGO enrichment of GPR37L1 co-IP proteins was conducted using the web-based server. The mouse reference proteome was used as background. Venn diagrams showing overlap between VCAM1 and GPR37L1 co-IP proteins were generated with the VennDiagram package (v1.7.3). Volcano plots were produced with the EnhancedVolcano package (v1.24.0) and all other plots were generated with ggplot2 (v3.4.3).

### In vitro synapse quantification assay

#### Commercial recombinant proteins

Upon arrival, lyophilized recombinant mouse VCAM1-Fc (R&D Systems, 643-VM) was briefly centrifuged and reconstituted in sterile Dulbecco’s phosphate-buffered saline (DPBS). Protein concentration was determined using the Pierce BCA protein assay kit (Thermo Scientific, 23227) according to the manufacturer’s instructions. Aliquots were prepared and snap-frozen in isopentane chilled at -60 to -70°C on dry ice, then stored at -80°C. Control Fc protein (ChromPure Human IgG Fc fragment, Jackson ImmunoResearch, 009-000-008) was stored at 4°C upon arrival.

#### Western blot quality control of recombinant proteins

For quality assessment, 2 µg of mouse VCAM1-Fc protein and Fc were mixed with 5X SDS loading buffer brought to volume with distilled water, and boiled for 10 min at 70 °C. Proteins were separated on a 4-20% precast polyacrylamide gel (Bio-Rad, 4561094) at 100V for ∼1.5 h and transferred to a 0.2 µm nitrocellulose membrane using a Trans-Blot Turbo Transfer System (Bio-Rad, 1704150EDU) under the mixed-molecular weight program. Membranes were stained with the Revert Total Protein Stain Kit (Li-COR, 926-11021) according to the manufacturer’s protocol and imaged on an Amersham ImageQuant 800 (Cytiva).

#### Primary hippocampal neuronal cultures and treatment

Primary hippocampal neurons were cultured from E18 to E19 dissected C57BL/6J WT embryos as previously described^74^. Briefly, dissociated neurons were plated on poly-D-lysine (PDL, Millipore) and laminin (Sigma-Aldrich) coated glass coverslips at a density of 30,000 cells per coverslip in MEM-horse serum medium. After 2 h, once neurons had adhered, the coverslips were flipped over an astroglial feeder layer in 12-well plates containing Neurobasal medium (Invitrogen) supplemented with B27 to establish sandwich co-cultures. Cultures were maintained at 37°C in a humidified 5% CO2 incubator. To inhibit glial proliferation, 10 mM glia inhibitor (5-fluoro-2-deoxyuridine, Sigma-Aldrich) was added at DIV3. Cultures were treated at DIV8 and DIV11 with 2 µg/mL recombinant mouse VCAM1-Fc (R&D Systems, 643-VM) or Fc control proteins (Jackson ImmunoResearch, 009-000-008). At DIV14, neurons were fixed in 4% paraformaldehyde (PFA)/4% sucrose in PBS for 10 mins and processed for ICC.

#### Immunocytochemistry (ICC)

Following fixation, coverslips were washed 3 x 5 min in PBS, permeabilized with 0.5% Triton X-100 in PBS for 10 min at RT in the dark. Afterwards, coverslips were washed for 3 x 5 min with PBS and neurons were blocked for 30 min with 10% BSA in PBS (2 µm-filtered). Primary antibodies (**Table S2**) were diluted in 3% BSA in PBS (2 µm-filtered) and incubated overnight at 4°C. The next day, coverslips were washed 3 x 5 min in PBS and incubated with fluorophore-secondary antibodies in 3% BSA in PBS (**Table S3**) for 1 at RT. Finally, neurons were washed 3 x 5 min in PBS and coverslips were mounted on microscope slides with Mowiol mounting medium. Importantly, antibodies for VGLUT1 and PSD95 were distinct from those used in the CRISPR/Cas9 KO in vivo experiments.

#### Image acquisition

Images were acquired using a Zeiss LSM880 confocal laser scanning microscope with a 63X oil objective (NA 1.4). All imaging was performed in confocal mode under identical acquisition settings across experimental conditions, to enable quantitative comparison. For each field of view (FOV), a Z-stack of four optical sections and a 2 × 2 tile scan were acquired to ensure full coverage of the region of interest. Images were captured at 1944 × 1944 pixels with a pixel size of 0.13 µm × 0.13 µm, a total z-stack thickness of 1.09 µm, and 16-bit depth. Five FOVs were acquired per coverslip, and 9 to 11 FOVs per condition in a single experiment. To minimize selection bias, only MAP2 staining was used for the initial localization of neuronal somas and dendrites, ensuring that at least two neuronal somas were located within proximity of each other before proceeding to acquisition. Pre- and postsynaptic channels were not consulted during field selection. For the VGLUT1-PSD95 dataset, image acquisition was performed blind to experimental condition. For the VGAT-GEPH dataset, image acquisition was not blinded.

#### Semi-automated image processing

In vitro excitatory (VGLUT1+PSD95+) and inhibitory (VGAT+GEPH+) synapses were analyzed using Fiji (ImageJ, NIH) with custom-written macros to standardize processing across experiments. For each experiment, optimal thresholds were determined manually per channel (pre, MAP2, post) after splitting and maximum intensity projections z-stacks. Mean thresholds were calculated from both control and treatment conditions and then applied uniformly to all images from both conditions. To remove non-specific somatic staining, neuronal soma regions were outlined manually and inside information was cleared. Binary masks were generated for all channels, and the pre- and postsynaptic channels were analyzed using the “Analyze Particles” function to quantify the synaptic puncta. MAP2+ dendritic arbors were skeletonized, and total branch length was extracted using the hiPNAT plugin. After each run, all open images and tables were closed automatically before proceeding to the next image. Few images were excluded during image processing owing to poor signal quality (two FOVs for VGLUT1-PSD95 dataset and two FOVs for the VGAT-GEPH dataset). All data (puncta counts, particle size, dendritic branch length) were manually compiled in Excel for downstream statistical analysis.

#### Downstream data analysis

The processed data in *XLSX* format were loaded into R (v4.5.0) running in a custom Docker development container (v28.1.1^67^). Raw data were imported from Excel files containing synaptic puncta counts, puncta sizes and total dendritic branch length measured per FOV. Across all experiments, control (Fc) mean was computed per metric and used to normalize corresponding Fc and VCAM1-Fc values. Normalized values were log2 transformed (log2 [1 + x]) to stabilize variance and accommodate zero values. Linear mixed-effects models were fitted using the lmer function from lme4 (v1.1-37) with treatment as a fixed effect and experiment as a random intercept to account for inter-experimental variability (∼experiment + condition). Separate models were generated for total, pre-, and postsynaptic puncta densities and puncta sizes. Model fit was assessed through visual inspection of residual plots and Q-Q plots to verify assumptions of normality and homoscedasticity. Estimated marginal means and pairwise contrasts between treatments were calculated using emmeans (v1.11.1). Pairwise differences between treatment levels were assessed using Tukey-corrected pairwise contrasts. Plots were generated with ggplot2 (v3.5.2) and ggbeeswarm (v0.7.2) showing individual FOVs that were normalized to Fc but not log2 transformed and per experiment means.

---

**Fig. S1.**
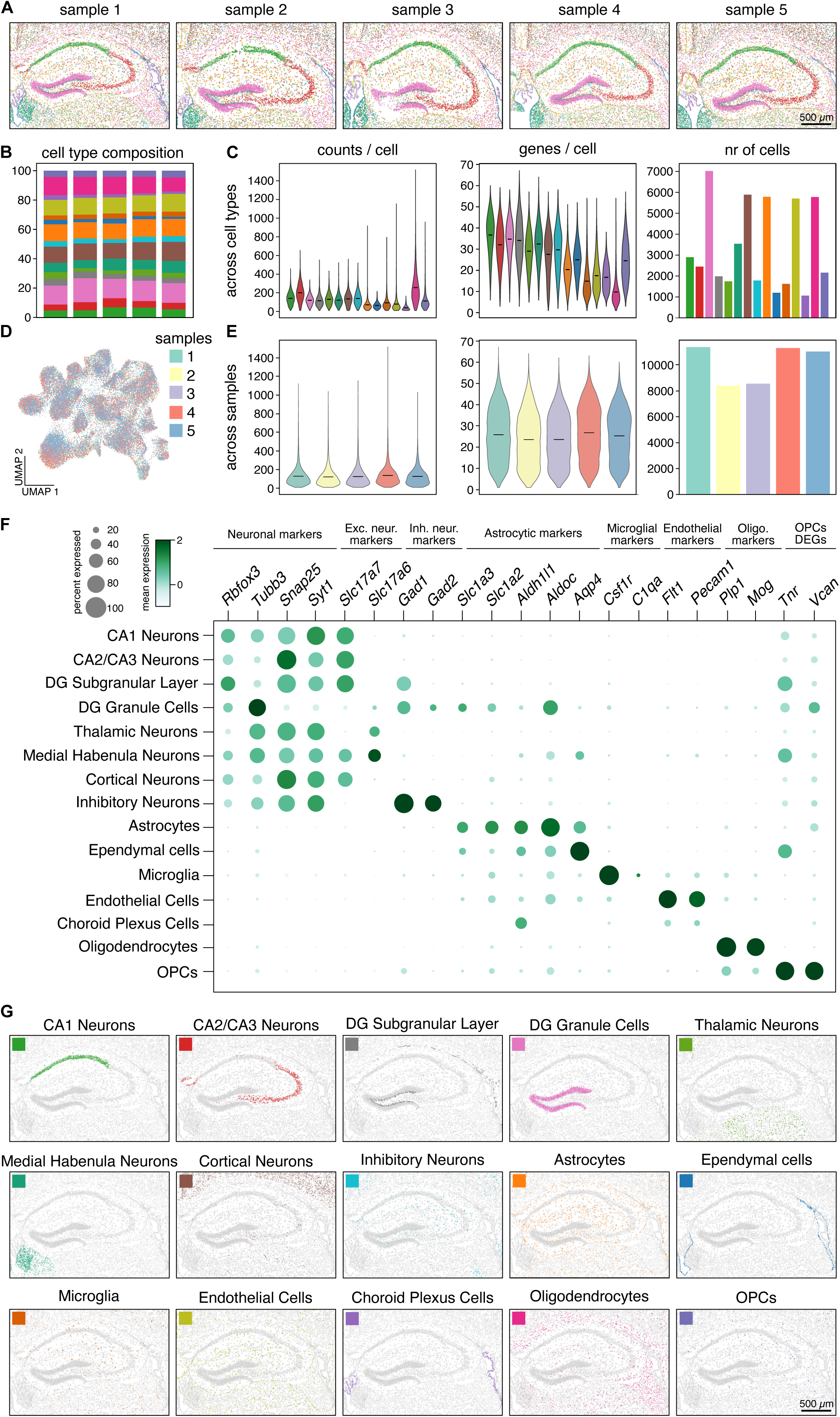
Quality control of targeted single-cell spatial transcriptomics. (A) Spatial plots showing the distribution of the 15 identified cell types across all samples. (B) Cell type composition across the samples, shown as percentages. (C) Violin and bar plots showing the number of counts per cell, number of genes per cell and number of cells for each of the identified cell types. (D) UMAP plot showing effective batch correction and integration across samples. (E) Violin and bar plots showing the number of counts per cell, number of genes per cell and number of cells across samples. (F) Dot plot showing cell type-specific genes across the 15 identified cell types from the batch-corrected dataset (negative expression values are possible because of the batch correction). OPCs were identified based on enriched expression of *Tnr* and *Vcan*. (G) Spatial plots showing each of the 15 identified cell types separately in one representative sample (sample 1).

**Fig. S2.**
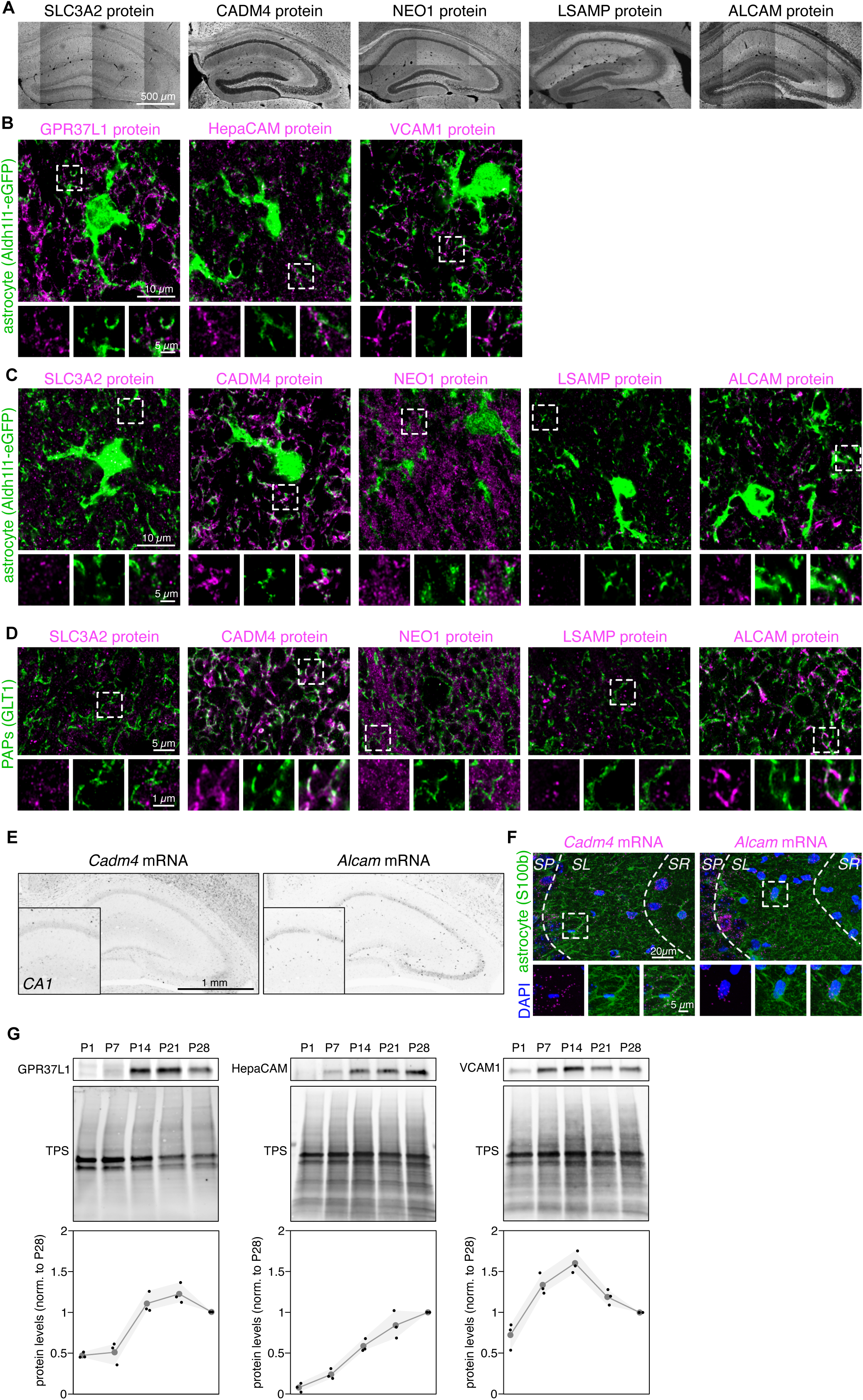
Additional profiling of GPR37L1, HepaCAM and VCAM1 and of non-high confidence astrocytic CSP candidates SLC3A2, CADM4, NEO1, LSAMP and ALCAM. (A) Low magnification single-plane immunostaining images of SLC3A2, CADM4, NEO1, LSAMP and ALCAM in the hippocampus of WT mice at P28. Representative of two independent experiments. (B) High magnification single-plane immunostaining images of GPR37L1, HepaCAM and VCAM1 in the CA3 SL region of Aldh1l1-eGFP mice at P28. Insets highlight colocalization with GFP. Representative of two independent experiments. (C) High magnification single-plane immunostaining images of SLC3A2, CADM4, NEO1, LSAMP and ALCAM in the CA3 SL region of Aldh1l1-eGFP mice at P28. Insets highlight full (CADM4), partial (SLC3A2, ALCAM), or no colocalization (NEO1, LSAMP) with GFP. Representative of two independent experiments. (D) High magnification single-plane immunostaining images of SLC3A2, CADM4, NEO1, LSAMP and ALCAM with the peri-synaptic astrocytic process marker GLT1 in the CA3 stratum lucidum (SL) of WT mice at P28. Insets highlight full (CADM4), partial (ALCAM) and no colocalization (SLC3A2, NEO1, LSAMP) with GLT1. Representative of two independent experiments. (E) Low magnification single-plane smFISH images of *Cadm4* and *Alcam* in the hippocampus of WT mice at P28. Representative of two independent experiments. (F) Representative smFISH z-stack images of *Cadm4* and *Alcam* in the CA3 region of WT mice at P28. Insets highlight expression in astrocytes labeled with S100β within the CA3 SL. Hippocampal layers were delineated by VGLUT1 co-staining (not shown). Representative of two independent experiments. (G) Western blot analysis of hippocampal lysates from littermate mice at across postnatal development, probed for GPR37L1, HepaCAM and VCAM. Protein levels were first normalized to total protein levels (TPS) and then to P28 levels. n = 3 animals per timepoint; mean ± 95% confidence interval.

**Fig. S3.**
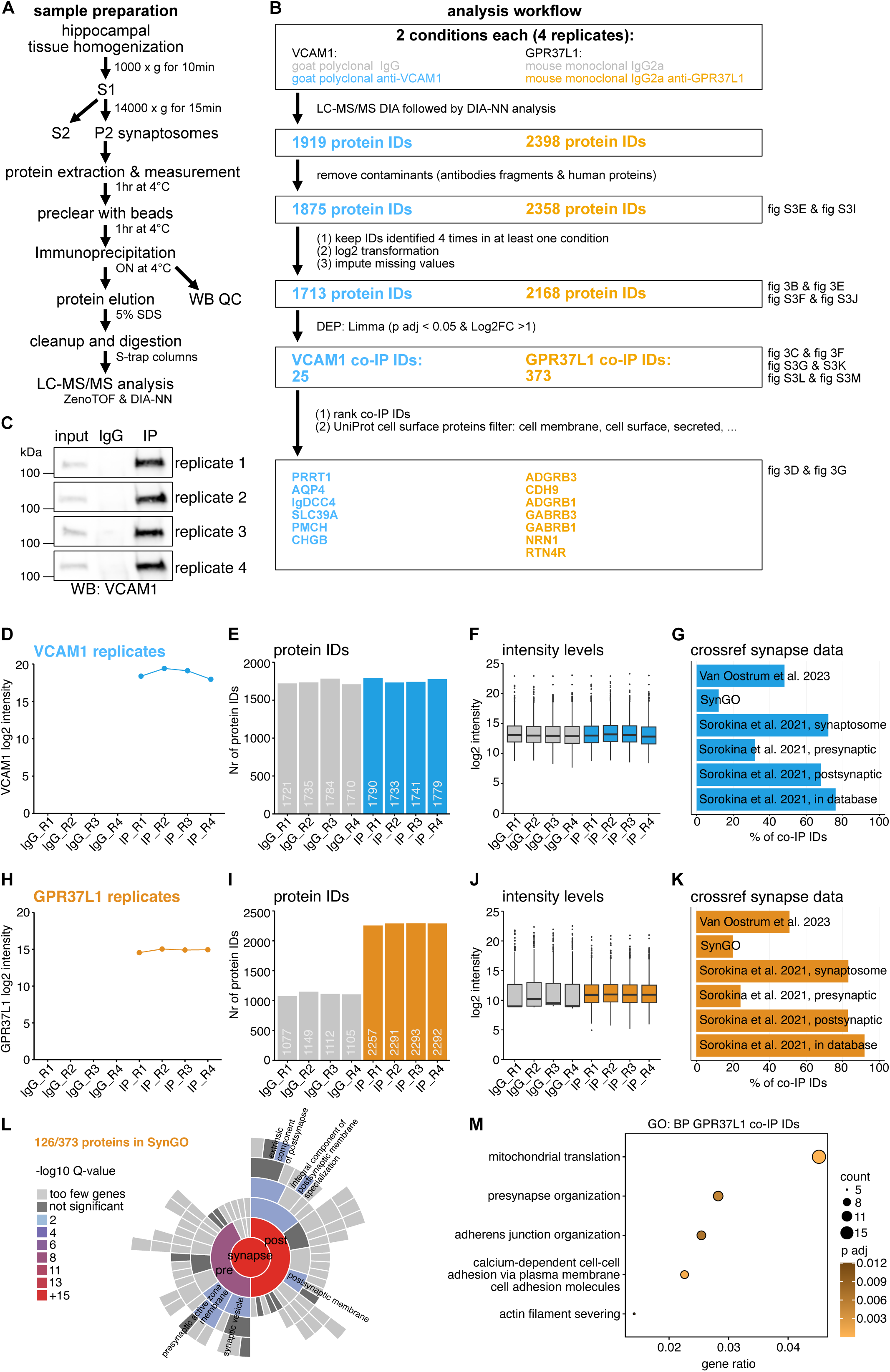
Methodology and quality control of synaptosomal VCAM1 and GPR37L1 co-immunoprecipitation proteomics. (A) Overview of sample preparation and synaptosomal co-IP workflow. (B) Computational analysis workflow for data processing and DEP identification (C) Western blot quality control of VCAM1 IP across four replicates used for MS-based proteomics. (D) VCAM1 log_2_ intensities per sample. (E) Number of unique proteins identified across VCAM1 replicates (F) Sample-level intensity distributions across VCAM1 replicates. (G) Cross reference of VCAM1 co-IP proteins with synaptic proteome databases and SynGO. (H) GPR37L1 log_2_ intensities per sample. (I) Number of unique proteins identified across GPR37L1 replicates (J) Sample-level intensity distributions across GPR37L1 replicates. (K) Cross reference of GPR37L1 co-IP proteins with synaptic proteome databases and SynGO. (L) SynGO analysis of GPR37L1 co-IP proteins. (M) Top 5 GO:BP terms of GPR37L1 co-IP proteins.

**Fig. S4.**
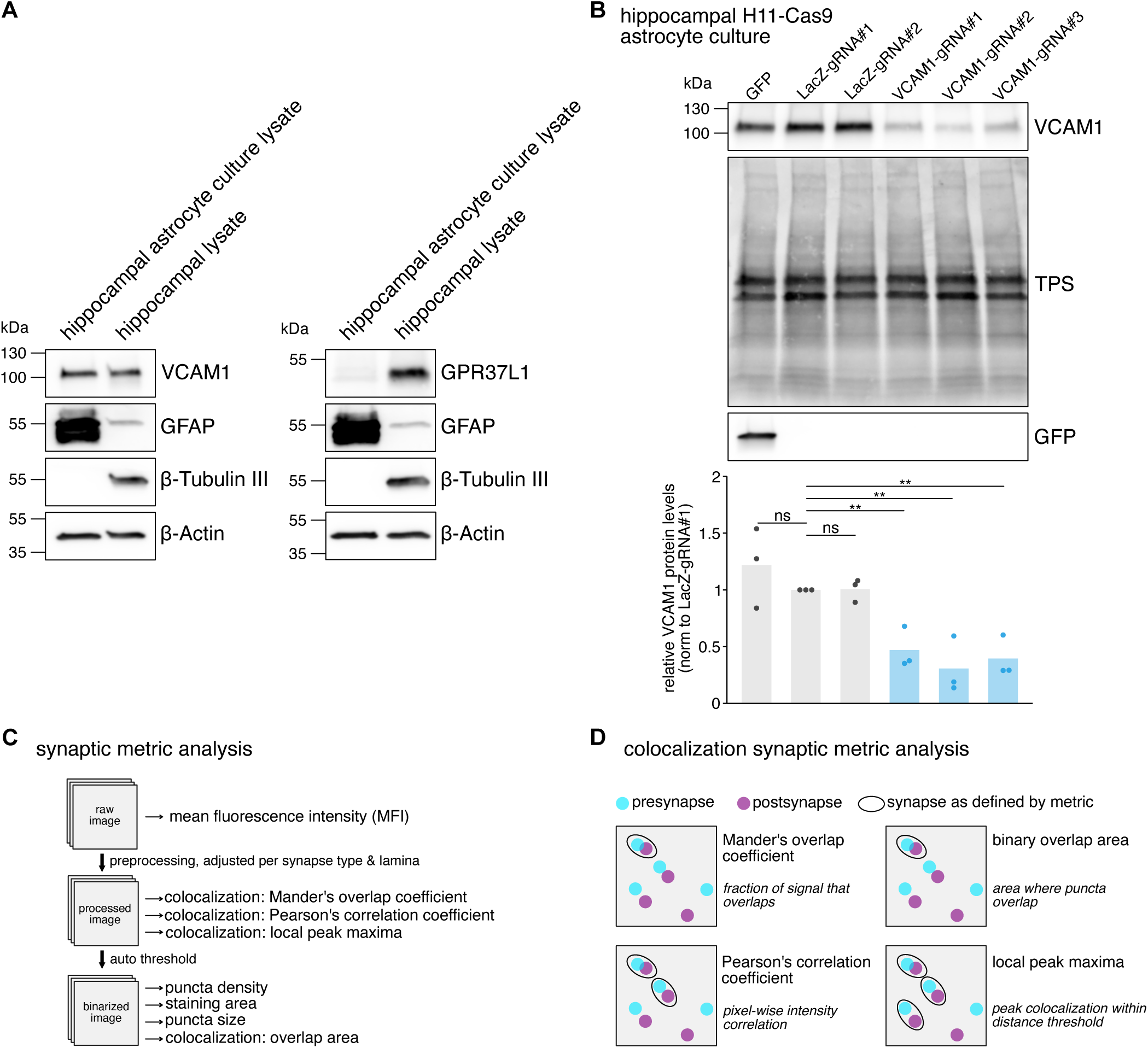
Quality control of gRNA targeting and synaptic metric illustrations. (A) Western blot analysis of hippocampal astrocyte culture lysates (DIV14-15) and whole hippocampal lysates (P28) from wild-type mice, probed for VCAM1 or GPR37L1 together with astrocytic marker GFAP, neuronal marker β-tubulin III and loading control β-actin. Representative of 2 independent experiments. TPS, total protein stain. (B) Western blot analysis and quantification of hippocampal H11-Cas9 astrocyte culture lysates that were infected with AAVs encoding GFP (infection control), two LacZ control gRNAs, and three independent VCAM1-targeting gRNAs. Quantification shows relative VCAM1 protein levels normalized first to total protein and subsequently to LacZ-gRNA#1. Data are from n = 3 independent experiments. Statistical significance was assessed by ANOVA with gRNA and experiment as factors, followed by Dunn’s post hoc test comparing each gRNA condition to the LacZ-gRNA#1 control with Benjamini-Hochberg correction for multiple comparisons. P values: ns non-significant, *p<0.05, **p<0.01. TPS, total protein stain. (C) Overview of synaptic metric analysis per FOV during image processing and analysis. (D) Visual explanation of the four synapse colocalization metrics analyzed as readout.

**Fig. S5.**
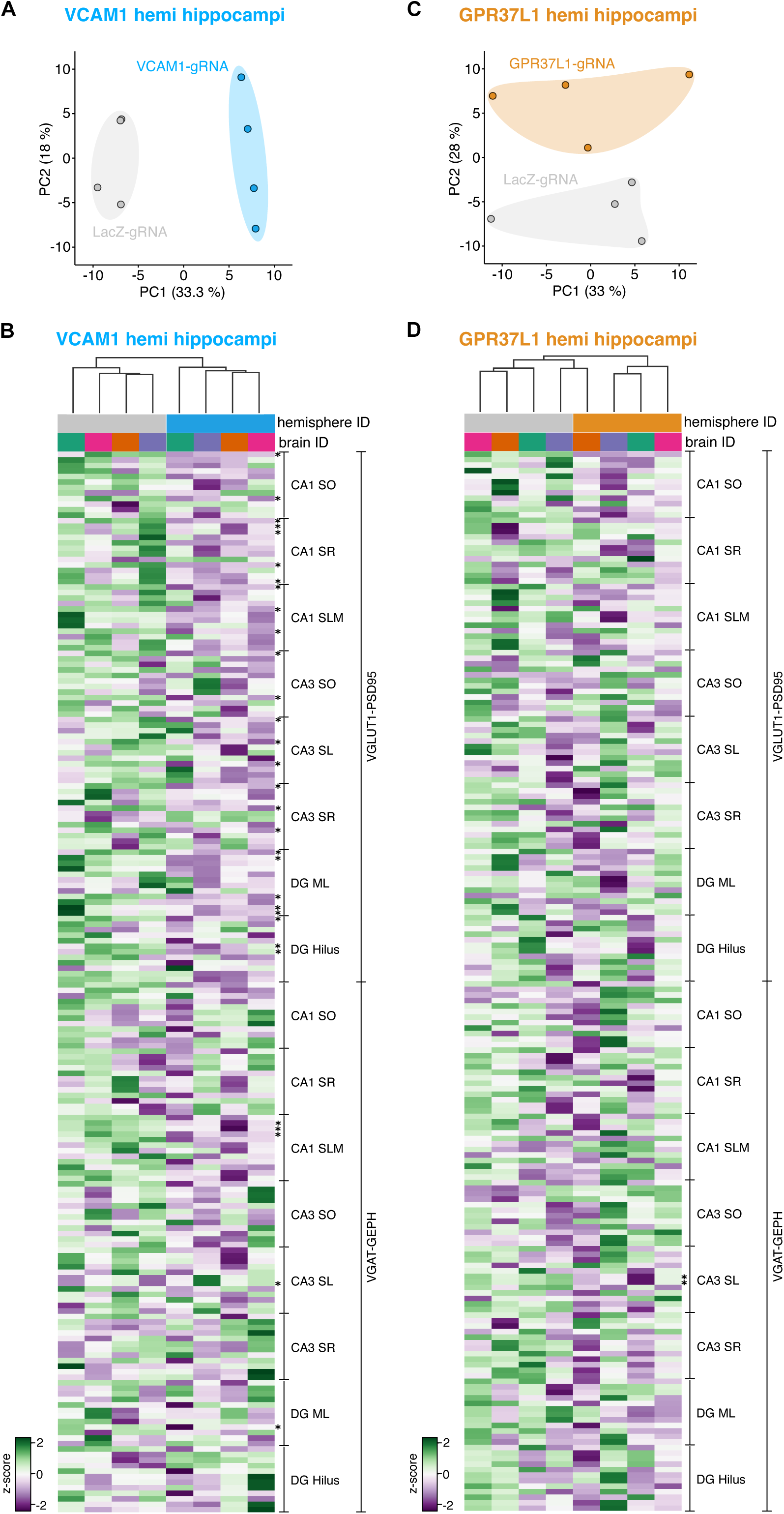
All synaptic metric quantifications across hippocampal hemispheres, synapse types and laminae. (A) PCA of synaptic metrics from LacZ-gRNA- and VCAM1-gRNA-injected hemispheres on residuals after linear mixed modeling showing clear separation by condition (n = 4 biological replicates). (B) Heatmap of all scaled synaptic metrics across the LacZ-gRNA- and VCAM1-gRNA-injected hemispheres and brains. Synaptic metrics were ordered according to Fig. 4G and metrics that were statistically significantly altered are highlighted with asterirks. (C) PCA of synaptic metrics of LacZ-gRNA- and GPR37L1-gRNA-injected hemispheres on residuals after linear mixed modeling (n = 4 biological replicates). (D) Heatmap of all scaled synaptic metrics across the LacZ-gRNA- and GPR37L1-gRNA-injected hemispheres and brains. Synaptic metrics were ordered according to Fig. 4H and metrics that that were statistically significantly altered are highlighted with asterirks.

**Fig. S6.**
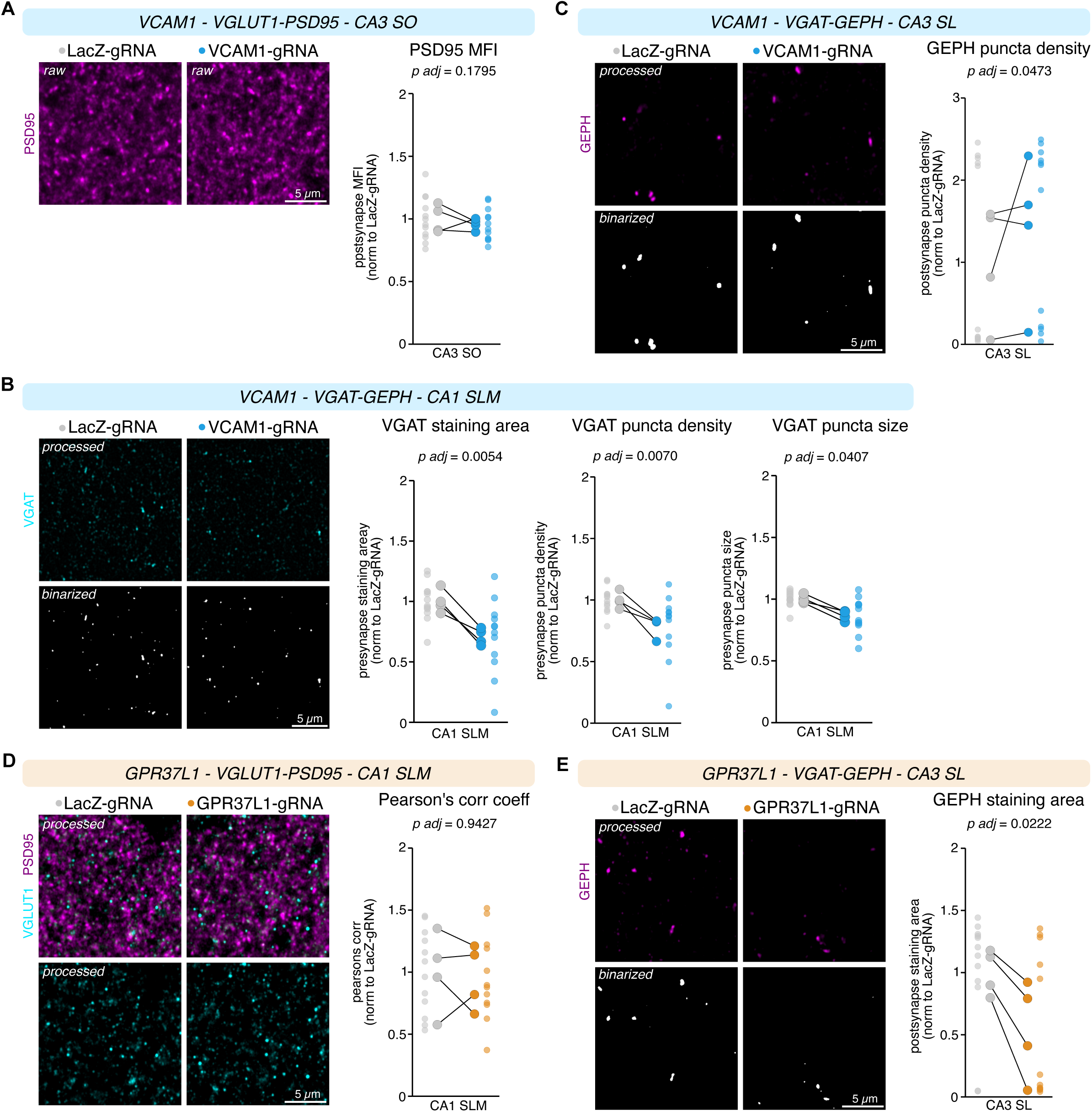
Additional representative examples of synaptic metrics. (A) Representative raw Airyscan images of VGLUT1-PSD95 in CA3 SO from LacZ-gRNA-and VCAM1-gRNA-injected hemispheres, with quantification of the PSD95 MFI normalized to the LacZ-gRNA hemisphere. Each small data point represents a single FOV; larger linked data points represent hemisphere means. Statistical significance was assessed using a linear mixed effects model as in Fig. 3G, followed by Tukey’s post hoc tests and FDR corrected significance. (B) Representative processed and binarized Airyscan images of VGAT in CA1 SLM from LacZ-gRNA- and VCAM1-gRNA-injected hemispheres, with quantification of presynaptic puncta density normalized to LacZ-gRNA. Same sampling, plotting and analyses as in (A). (C) Representative processed and binarized Airyscan images of GEPH in CA3 SL from LacZ-gRNA- and VCAM1-gRNA-injected hemispheres, with quantification of GEPH staining area normalized to LacZ-gRNA. Same sampling, plotting and analyses as in (A). (D) Representative processed Airyscan images of VGLUT1-PSD95 in CA3 SLM from LacZ-gRNA- and GPR37L1-gRNA-injected hemispheres, with quantification of VGLUT1-PSD95 Pearson’s correlation normalized to LacZ-gRNA. Same sampling, plotting and analyses as in (A). (E) Representative processed and binarized Airyscan images of GEPH in CA3 SL from LacZ-gRNA- and GPR37L1-gRNA-injected hemispheres, with quantification of GEPH staining area normalized to LacZ-gRNA. Same sampling, plotting and analyses as in (A).

**Fig. S7.**
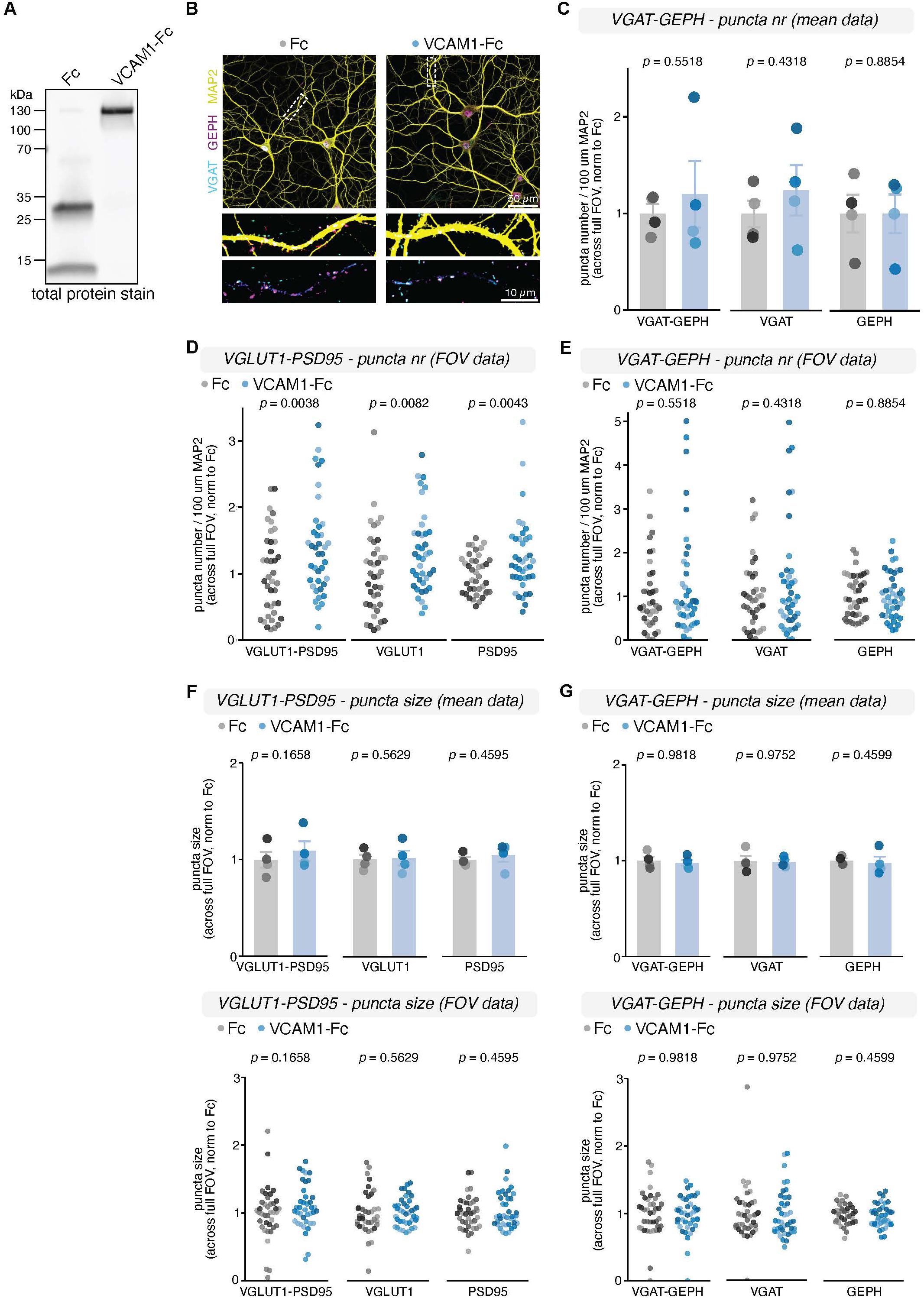
Recombinant VCAM1 increases excitatory but not inhibitory hippocampal synapse density *in vitro*. (A) Western blot quality control of the recombinant proteins Fc and VCAM1-Fc used to treat hippocampal neurons, visualized using a total protein stain. (B) Representative images of hippocampal neurons treated with recombinant protein from DIV8 to DIV14. Fc or VCAM1–Fc (2 µg/mL) was added at DIV8 and replenished at DIV11. Neurons were fixed at DIV14 and stained for VGAT (cyan), GEPH (magenta), and MAP2 (yellow). Insets show the merge followed by the isolated VGAT-GEPH signal. (C) Quantification of inhibitory synaptic puncta density along MAP2+ dendrites. VGAT+GEPH+ colocalized puncta, as well as VGAT+ and GEPH+ puncta separately, were measured per 100 μm dendrite and normalized to Fc. Same sampling, plotting and analyses as in Fig. 4J. (D) Plots from Fig. 4J showing distribution of the analyzed FOVs. Each dot represents a single FOV and is color-shaded according to experiment. (E) Plots from (C) showing distribution of the analyzed FOVs. Each dot represents a single FOV and is color-shaded according to experiment. (F) Quantification of excitatory synaptic puncta size of VGLUT1+PSD95+ colocalized puncta, as well as VGLUT1+ and PSD95+ puncta separately. Each FOV contained two neuronal somas for standardized sampling; 9-11 FOVs per condition were analyzed in N = 4 independent experiments. Each data point represents one experiment (above panel), and one FOV (below panel), color-shaded by experiment. A linear mixed-effects model was used to account for multiple FOVs per experiment. (G) Quantification of inhibitory synaptic puncta size of VGAT+GEPH+ colocalized puncta, as well as VGAT+ and GEPH+ puncta separately. Same sampling, plotting and analyses as in (F).

**Table S1.** Tested antibodies in IHC and WB for CSP candidate profiling.

| CSP candidate | Species | Clonality | Source | Ref | IHC | WB | Validation |
| --- | --- | --- | --- | --- | --- | --- | --- |
| GPR37L1 | Mouse | Mono IgG1 | MabTech | GPR37L1-2 | ✓ | - | IHC KO:<br>https://pubmed.ncbi.nlm.nih.gov/24062445/ |
|  | Rabbit | Poly | MabTech | GPR37L1-3 |  | - |  |
|  | Rabbit | Poly | Invitrogen | PA5-106838 | - | - |  |
|  | Rabbit | Poly | Novus | NLS405 | - | - |  |
|  | Rabbit | Mono | Cell Signaling | #93782 |  | - |  |
|  | Mouse | Mono IgG2a | R&D Systems | MAB4449 | ✓ | ✓ | IHC KO:<br>this paper |
| HepaCAM | Mouse | mono IgG1 | R&D Systems | MAB4108 | ✓ | ✓ | IP KO:<br>https://pubmed.ncbi.nlm.nih.gov/34171291/<br>IHC KO:<br>https://pubmed.ncbi.nlm.nih.gov/25382142/ |
| PLXNB1 | Goat | Poly | R&D Systems | AF3749 |  | - | WB KO:<br>https://pubmed.ncbi.nlm.nih.gov/26579598/ |
|  | Rabbit | Poly | Merck | HPA040586 | - | - |  |
|  | Mouse | Mono IgG1 | R&D Systems | MAB3749 | - | - |  |
|  | Rabbit | Poly | Abcam | ab90087 | - | - | WB KO:<br>https://pubmed.ncbi.nlm.nih.gov/29242274/ |
| VCAM1 | Goat | Poly | R&D Systems | AF643 | ✓ | ✓ | IHC KO & WB KO:<br>this paper |
|  | Rabbit | Mono | Abcam | ab134047 | - | - | WB KD:<br>https://pubmed.ncbi.nlm.nih.gov/29670110/ |
|  | Rat | Mono | BD Biosciences | 550547 | - | - | IHC KO:<br>https://pubmed.ncbi.nlm.nih.gov/31907535/ |
|  | Rat | Mono | Millipore | CBL1300 | - | - |  |
| LSAMP | Mouse | Mono IgG2a | DSHB | 2G9-c | ✓ | ✓ |  |
| NEO1 | Goat | Poly | R&D Systems | AF1079 | ✓ | ✓ | WB KO & IHC KO:<br>https://pubmed.ncbi.nlm.nih.gov/27143755/ |
| ALCAM | Goat | Poly | R&D Systems | AF1172 | ✓ | ✓ | WB KO:<br>https://pubmed.ncbi.nlm.nih.gov/28069965/<br>IHC KO:<br>https://pubmed.ncbi.nlm.nih.gov/20016077/ |
| SynCAM4 | Mouse | Mono IgG1 | NeuroMab | 75-247 | ✓ | ✓ | WB KO:<br>https://pubmed.ncbi.nlm.nih.gov/23700466/ |
| SLC3A2 | Rabbit | Poly | Merck | HPA017980 | ✓ | ✓ | WB KD:<br>https://pubmed.ncbi.nlm.nih.gov/32093034/ |
|  | Rabbit | Poly | Abnova | H00006520 | ✓ | ✓ | WB KD:<br>https://pubmed.ncbi.nlm.nih.gov/23785200/ |

**Table S2.** Primary antibodies used for IHC, ICC, WB & IP.

| Target | Species | Clonality | Source | Ref | Application & dilution |
| --- | --- | --- | --- | --- | --- |
| VCAM1 | Goat | Poly | R&D Systems | AF643 | IHC & WB: 1/1000, IP: 10 µg/mL |
| Control IgG | Goat | Poly | R&D Systems | AB108-C | IP: 10 µg/mL |
| GPR37L1 | Mouse | Mono IgG2a | R&D Systems | MAB4449 | IHC & WB: 1/1000, IP: 10 µg/mL |
| Control IgG | Mouse | Mono IgG2a | R&D Systems | MAB0031 | IP: 10 µg/mL |
| MAP2 | Chicken | Poly | Encor | CPCA-MAP2 | ICC: 1/1000 |
| PSD95 | Mouse | Mono IgG1 | ThermoFischer | MA1-046 | ICC: 1/200, WB: 1/2000 |
| PSD95-AF647 | Camelid | Nanobody | Synaptic Systems/NanoTag | N3702-AF647-L | IHC: 1/500 |
| VGLUT1 | Rabbit | Poly | Synaptic Systems | 135 302 | ICC: 1/1000 |
| VGLUT1 | Chicken | Poly | Synaptic Systems | 135 316 | IHC: 1/500 |
| VGLUT1 | Guinea Pig | Poly | Synaptic Systems | 135 318 | smFISH-IHC: 1/500 |
| Gephyrin | Rabbit | Mono | Synaptic Systems | 147 008 | ICC & IHC: 1/500 |
| VGAT | Guinea Pig | Poly | Synaptic Systems | 131 004 | ICC: 1/1000, IHC: 1/500 |
| GFAP | Guinea Pig | Poly | Synaptic Systems | 173004 | WB: 1/500 |
| β-III Tubulin | Rabbit | Poly | Abcam | ab18207 | WB: 1/10.000 |
| β-Actin | Mouse | Mono IgG2a | Sigma | A2228 | WB: 1/1000 |
| S100b | Rabbit | Mono | Abcam | ab52642 | smFISH-IHC: 1/1000 |
| GFP | Chicken | Poly | Aves | GFP-1010 | IHC: 1/1000 |
| GLT1 | Guinea Pig | Poly | Millipore | AB1783 | IHC: 1/500 |
| HepaCAM | Mouse | Mono IgG1 | R&D Systems | MAB4108 | WB: 1/1000, IHC: 1/100 |
| Ezrin | Rabbit | Poly | Cell Signaling | 3145 | WB: 1/2000 |
| SYP | Mouse | Mono IgG1 | Sigma | S5768 | WB: 1/2000 |
| SLC3A2 | Rabbit | Poly | Abnova | H00006520 | IHC: 1/100 |
| CADM4 | Mouse | Mono IgG1 | NeuroMab | 75-247 | IHC: 1/100 |
| NEO1 | Goat | Poly | R&D Systems | AF1079 | IHC: 1/200 |
| LSAMP | Mouse | Mono IgG2a | DSHB | 2G9-c | IHC: 1/1000 |
| ALCAM | Goat | Poly | R&D Systems | AF1172 | IHC: 1/200 |

**Table S3.** Secondary antibodies used for IHC, ICC, WB & IP.

| Secondary antibody | Primary target(s) | Source | Ref | Application & dilution |
| --- | --- | --- | --- | --- |
| Rabbit anti-Goat HRP | VCAM1 | ThermoFischer | 81-1620 | WB: 1/10.000 |
| Donkey anti-Goat A555 | VCAM1 | ThermoFischer | A21432 | IHC: 1/1000 |
| Goat anti-Mouse IgG2a HRP | GPR37L1, β-Actin | ThermoFischer | A10685 | WB: 1/1000 |
| Goat anti-Mouse IgG2a A555 | GPR37L1 | ThermoFischer | A21137 | IHC: 1/1000 |
| Goat anti-Chicken A555 | MAP2 | ThermoFischer | A78949 | ICC: 1/1000 |
| Goat anti-Mouse IgG1 A647 | PSD95, CADM4 | ThermoFischer | A21240 | ICC: 1/1000 |
| Donkey anti-Rabbit A488 | VGLUT1, S100b | ThermoFischer | A21206 | ICC & smFISH-IHC: 1/1000, |
| Donkey anti-Chicken A488 | VGLUT1 | ThermoFischer | A78948 | IHC: 1/1000 |
| Donkey anti-Rabbit A647 | Gephyrin, SLC3A2 | ThermoFischer | A31573 | ICC & IHC: 1/1000 |
| Donkey anti-Guinea Pig A488 | VGAT | Jackson ImmunoResearch | 706-545-148 | ICC & IHC: 1/250 |
| Donkey anti-Guinea Pig A647 | VGLUT1 | Jackson ImmunoResearch | 706-605-148 | smFISH-IHC: 1/250 |
| Goat anti-Guinea Pig HRP | GFAP | ThermoFischer | A18769 | WB: 1/10.000 |
| Goat anti-Rabbit HRP | β-III Tubulin, Ezrin | ThermoFischer | 65-6120 | WB: 1/5000 |
| Goat anti-Mouse IgG1 HRP | HepaCAM, PSD95, SYP | ThermoFischer | A10551 | WB: 1/1000 |
| Donkey anti-Goat A488 | NEO1 | ThermoFischer | A11055 | IHC: 1/1000 |
| Goat anti-Mouse IgG2a A647 | LSAMP | ThermoFischer | A21241 | IHC: 1/1000 |
| Donkey anti-Goat A647 | ALCAM | ThermoFischer | A21447 | IHC: 1/1000 |

**Table S4.** Probes used for smFISH-IHC (RNAscope).

| Target | Probe | Source | Ref | Dilution |
| --- | --- | --- | --- | --- |
| Gpr37l1 | Mm-Gpr37l1 | ACDBio | 319301 |  |
| Hepacam | Mm-Hepacam | ACDBio | 476101 |  |
| Vcam1 | Mm-Vcam1 | ACDBio | 438641 |  |
| Alcam | Mm-Alcam | ACDBio | 462061 |  |
| Cadm4 | Mm-Cadm4-C1 | ACDBio | 1167411-C1 |  |
| DapB negative control | Mm-DapB | ACDBio | 310043 |  |
| C1 | Opal Dye 570 | Akoya Biosciences | FP1488001KT | 1:1000 |

**Table S5.**
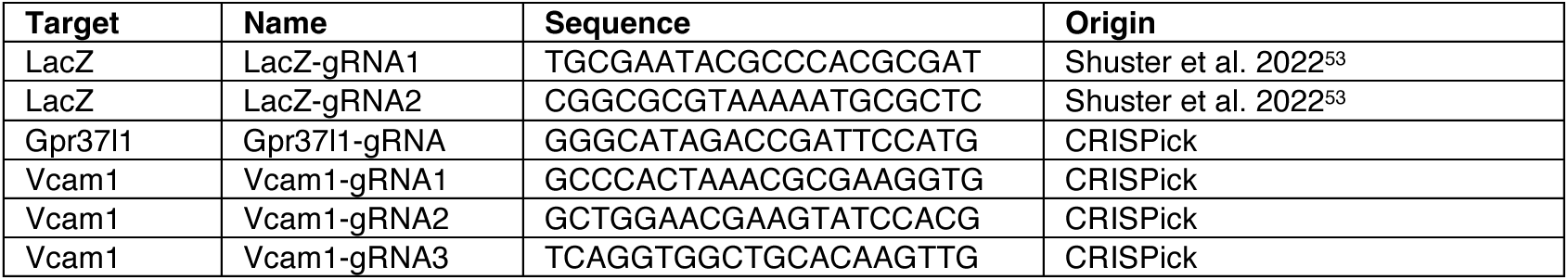
Guide RNA sequences for CRISPR/Cas9 KO.

**Table S6.**
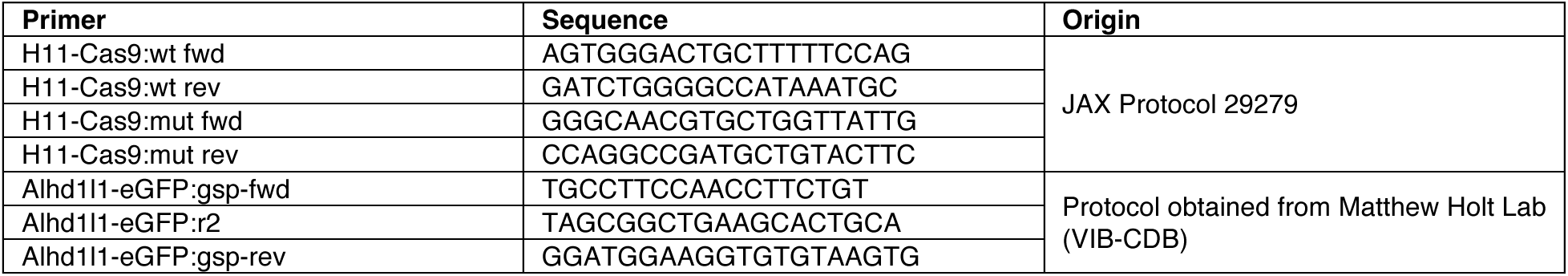
Genotyping PCR primers.

